# The transcriptomic and proteomic ramifications of segmental amplification

**DOI:** 10.1101/2023.11.21.568005

**Authors:** Ryan K. Fritts, Christopher C. Ebmeier, Shelley D. Copley

## Abstract

Gene amplification can drive adaptation by rapidly increasing the cellular dosage of critical gene products. Segmental amplifications often encompass large genomic regions surrounding the gene(s) under selection for higher dosage. Overexpression of co-amplified neighboring genes imposes a substantial metabolic burden. While compensatory mutations can decrease inappropriate overexpression of co-amplified genes, it takes time for such mutations to arise. The extent to which intrinsic regulatory mechanisms modulate expression of co-amplified genes in the immediate aftermath of segmental amplification is largely unknown. To address the collateral effects of segmental amplification, we evolved replicate cultures of an *Escherichia coli* mutant under conditions that select for higher dosage of an inefficient enzyme whose weak activity limits growth rate. Segmental amplifications encompassing the gene encoding the weak-link enzyme arose in all populations. Amplicons ranged in size (9 to 125 kb) and copy number (2 to 12 copies). We performed RNA-seq and label-free proteomics to quantify expression of amplified genes present at 2, 6, and 12 copies. mRNA expression generally scales with gene copy number, but protein expression scales less well with both gene copy number and mRNA expression. We characterize the molecular mechanisms underlying discrepancies between gene copy number and expression for several cases. We also show that segmental amplifications can have system-wide consequences by indirectly altering expression of non-amplified genes. Our findings indicate that the fitness benefit derived from segmental amplification depends on the combined effects of amplification size, gene content, and copy number as well as collateral effects on non-amplified genes.

**Significance Statement:** Gene amplification frequently drives rapid adaptation when organisms are challenged by harsh conditions. However, gene amplification rarely amplifies only the gene(s) under selection for higher dosage. Segmental amplifications often include many co-amplified neighbor genes. Because segmental amplifications can reach very high copy number, overexpression of co-amplified genes can impose an energetic burden and perturb physiology, yet this aspect of gene amplification has received little attention. Here, we performed transcriptomic and proteomic analysis of laboratory-evolved *Escherichia coli* strains containing large, high-copy-number amplifications. We found that mRNA levels, but often not protein levels, scale well with gene copy number. We identified amplified genes that exhibit discrepancies between gene copy number and mRNA/protein expression and examine the molecular mechanisms underlying these discrepancies.

## Introduction

Organisms frequently encounter new and/or changing environments. Successful adaptation to a new niche may require an increase in the expression of particularly important gene products. Gene amplification allows organisms to rapidly increase the cellular dosage of newly needed proteins. Gene amplification has been shown to be adaptive under various contexts for bacteria (1–10) and eukaryotes (11–18) and is implicated in the development and proliferation of cancer cells (19–23).

Beyond serving as a mechanism for rapidly increasing the dosage of a gene product, gene amplification can also be the starting point for evolution of new molecular functions. The Innovation-Amplification-Divergence (IAD) model describes an evolutionary scenario by which a gene encoding a new function can arise (2, 24) (Fig. 1A). Most, if not all, proteins have weak side activities, termed promiscuous activities, in addition to their native activities. If a promiscuous activity becomes important for fitness (innovation), natural selection for increased expression often leads to gene amplification.

**Fig. 1.**
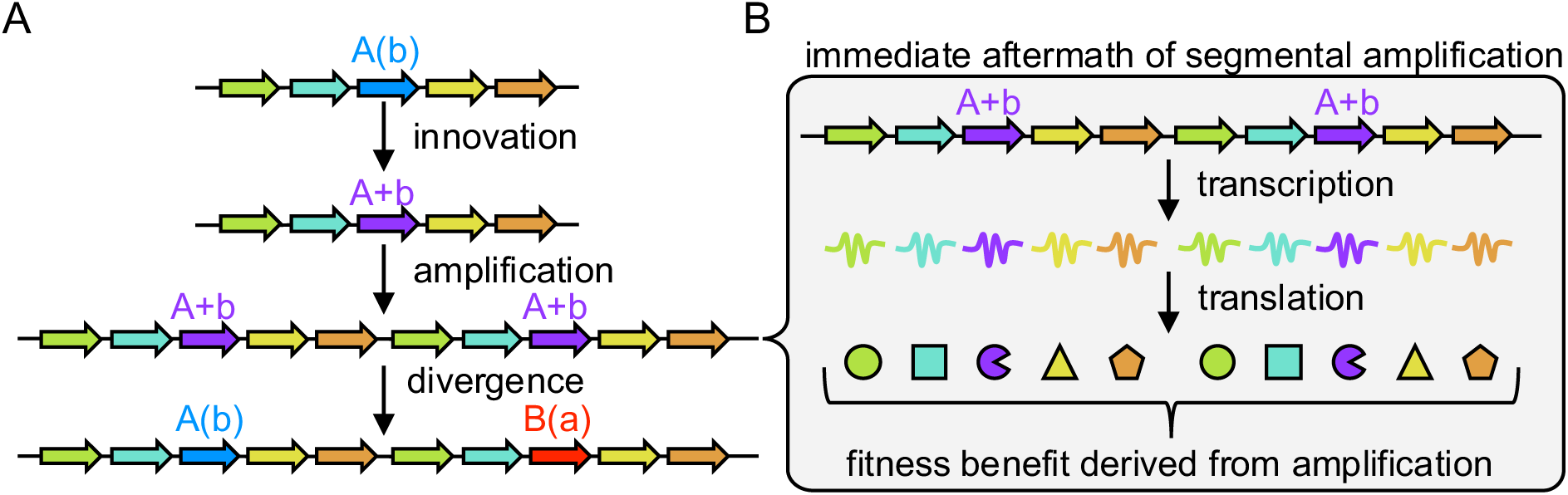
The expression and function of co-amplified neighboring genes can affect the fitness benefit derived from segmental amplification. (A) The Innovation-Amplification-Divergence (IAD) model. Capital letters, major function; lower-case letters, inefficient function; parentheses, promiscuous function. (B) The transcription and translation of all amplified genes determines the overall fitness benefit gained in the immediate aftermath of segmental amplification.

Following amplification, mutations in a new gene copy can improve the newly advantageous but weak activity while the ancestral function is maintained in another gene copy (divergence). This process has been responsible for the evolution of vast superfamilies of enzymes, transporters, transcriptional regulators, and signaling proteins (25–29).

Gene amplification does not neatly increase the copy number of a single gene under selection for higher expression. Rather, segmental amplifications often amplify large genomic regions containing dozens to hundreds of genes surrounding the gene(s) under selection for higher dosage. Segmental amplifications in bacteria and yeast often reach above 10 copies and/or span over 100 kb during adaptation to stresses such as starvation or antimicrobial exposure (4–10, 14, 18, 30, 31). Despite the evolutionary importance of gene amplification, little attention has been given to the consequences of co-amplifying neighboring genes (Fig. 1B). Co-amplification of neighboring genes would be expected to inappropriately increase levels of the encoded proteins and cause physiological perturbations and/or energetic burdens. These collateral effects could have important repercussions on population dynamics in communities of organisms that are competing to adapt in the face of an environmental challenge or opportunity.

Previous investigations of the levels of mRNAs and proteins produced from duplicated/amplified genes in aneuploid yeast strains with one or two extra copies of whole chromosomes (32–37) and in cancer cells (38, 39) have shown that mRNA and, to a lesser extent, protein levels often, but not always, scale with gene copy number. Alterations in expression of co-amplified genes in these settings may be due to either intrinsic regulatory mechanisms that counteract inappropriate gene expression or to compensatory mutations that arose over many generations. A large-scale examination of hundreds of natural aneuploid yeast isolates found that most proteins expressed from aneuploid chromosomes exhibited attenuated levels (37), a phenomenon termed dosage compensation. Because these stable aneuploidies present in natural yeasts are likely long-standing adaptations, there has been time for multiple compensatory mutations to arise to counteract the most detrimental consequences of aneuploidy and alter the proteome of these isolates. Another recent study assessing the expression of individually duplicated yeast genes similarly found that protein levels were attenuated in some cases, although the precise molecular mechanisms that dampened protein expression were usually not clear (40). The expression of high-copy-number co-amplified genes in the immediate aftermath of segmental amplification is largely unknown. Consequently, we lack a broad understanding of the intrinsic capacity of regulatory mechanisms to buffer against the collateral effects of segmental amplification. These regulatory mechanisms can operate at the transcriptional, posttranscriptional and posttranslational levels to attenuate expression. The level at which such regulation occurs affects the energetic burden incurred following segmental amplification. Repressing the transcription of a high-copy-number operon is obviously less energetically costly than degrading the encoded proteins following translation. However, the findings of several studies indicate that transcript levels generally correspond to gene copy number and that most dosage compensation happens at the posttranscriptional and posttranslational levels (17, 33–37, 41)

We have developed a model system in *Escherichia coli* to characterize the immediate ramifications of segmental amplification on the transcriptome and proteome. The *E. coli* enzymes ArgC and ProA are nonhomologous reductases that catalyze similar reactions in arginine and proline biosynthesis, respectively (Fig. 2). We previously deleted *argC* from the *E. coli* genome and selected for recruitment of another enzyme to replace the missing ArgC activity during growth on minimal medium (42). A strain with a point mutation in *proA*, which causes substitution of Glu383 to Ala in the active site of ProA, was able to restore arginine production. E383A ProA (ProA*) catalyzes both its native reaction in proline synthesis and the weak promiscuous activity required for arginine synthesis (Fig. 2). The poor activity of ProA* compromises growth in minimal media in which synthesis of proline and arginine is required for growth, leading to strong selective pressure for amplification of *proA*.* We previously observed segmental amplifications encompassing *proA*\* ranging in size from 5-165 kb and reaching copy numbers between 3-20 copies in experimentally evolved *E. coli* populations (4, 43).

**Fig. 2.**
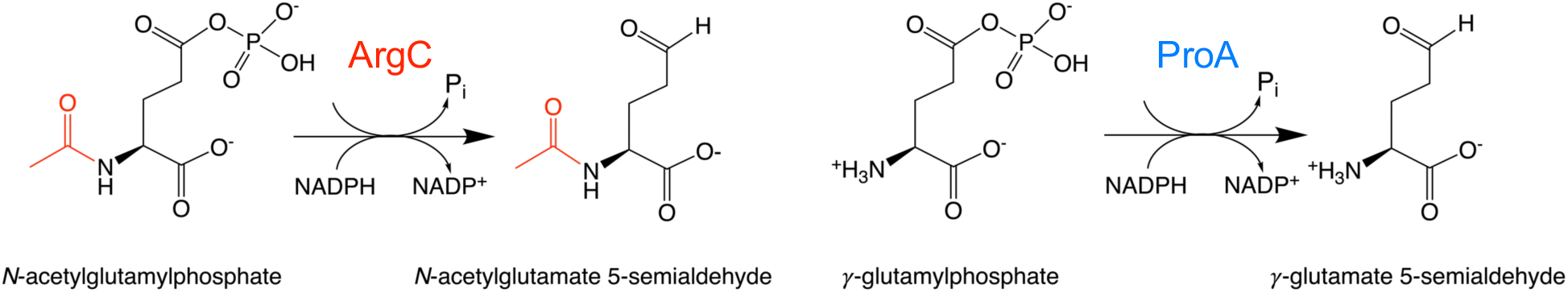
Reactions catalyzed by the reductases ArgC and ProA in *E. coli*. The acetyl group of *N*-acetylglutamylphosphate (red) is the only difference between the two substrates.

Many of the genes neighboring the *proBA*\* operon are pseudogenes or uncharacterized cryptic prophage genes. Thus, amplification of *proA\** in its native locus does not provide an ideal system for assessing the effects of co-amplification of genes near the gene under selection for higher copy number. For this work, we relocated the *proBA*\* operon to a new site neighboring a gene that we hypothesized might cause physiological perturbations if overexpressed following segmental amplification. We evolved eight replicate populations of this *E. coli* strain for 145-200 generations in a turbidostat. Whole genome sequencing revealed that all populations contained segmental amplifications of differing sizes and copy numbers. We examined the transcriptomes and proteomes of two populations, those with the highest copy number and largest amplifications, to determine the collateral effects of segmental amplification. The expression of many amplified genes scales with gene copy number at the mRNA and protein levels.

However, in several cases, mRNA and/or protein levels either exceed or fall short of the level expected based solely on gene copy number. We have examined the molecular mechanisms for several cases in which such discrepancies exist. We also found significant changes in levels of hundreds of non-amplified genes, demonstrating the profound and global impact of segmental amplification on cellular physiology. Through this survey of the system-wide consequences of segmental amplification, we identified a compensatory mutation that appears to prevent overexpression of unnecessary proteins, which may represent a broad class of adaptations to proteotoxic stress following gene amplification.

## Results and Discussion

### Segmental amplification increases growth rate of an *E. coli* strain with the *proBA\** operon relocated next to *rpoS*

We deleted the *proBA*\* operon from its native locus and relocated it next to *rpoS,* which encodes the alternative sigma factor RpoS/σ^S^ (also known as σ^38^) (Fig. 3A). We included the upstream region containing a C to T promoter mutation 45 bp upstream of *proB* that increases transcription of *proBA*\* (44). The *proBA*\* operon was oriented convergently to *rpoS* so that transcriptional read-through from *proBA*\* would not increase *rpoS* expression. A dual *rrnB* T1/T7 TE transcriptional terminator was placed immediately after *proA*\*. The alternative sigma factor RpoS serves as a global regulator by interacting with RNA polymerase to activate transcription of numerous genes important for redirecting cellular resources from growth toward survival (45, 46). Consequently, expression of RpoS is typically associated with a slowing of growth during stressful conditions or limited nutrient availability. Elevated expression of genes in the RpoS regulon confers improved resistance to starvation, high temperatures, low pH, oxidative stress and osmotic shock (45, 46). We hypothesized that overexpression of RpoS following co-amplification with *proA*\* might perturb cellular physiology because it would shift cells toward a transcriptional profile intended as a stress response to halt growth while the cells are growing exponentially with adequate nutrients.

**Fig. 3.**
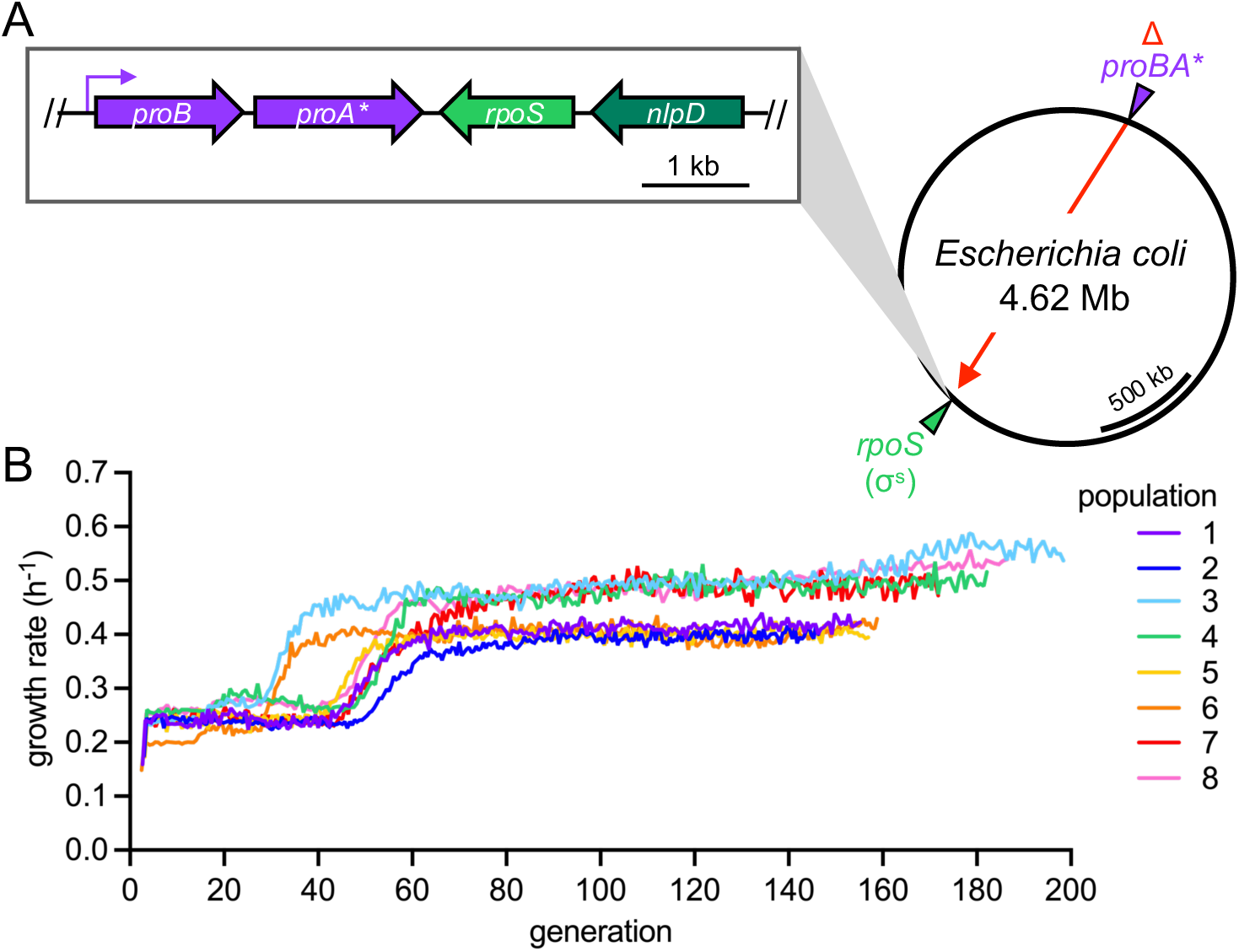
Evolution of an *E. coli* strain with the *proBA*\* operon relocated next to *rpoS*. (A) The *proBA*\* operon with a mutant constitutive promoter was deleted from its native locus and relocated directly next to the *rpoS* gene. (B) Growth rate increased during laboratory evolution of the *proBA*\*-*rpoS* relocation strain in minimal medium in a turbidostat.

We used a single founder colony of the Δ*argC proBA\** relocation strain to start eight replicate populations in a turbidostat. The cultures were grown in M9 minimal medium with glucose as the sole carbon source to impose strong selection for mutations that increase ProA* dosage. After 30-60 generations, we observed improvements in growth rate for all populations (Fig. 3B). Populations 3 and 8 showed an additional slight increase in growth rate around generation 180. We ended the experiment after 145-200 generations to limit the opportunity for compensatory mutations that might alleviate harmful effects of segmental amplification to arise. Similarly, we wanted to limit the chance for amplicon remodeling to remove unnecessary copies of amplified genes (43).

We sequenced genomic DNA (gDNA) isolated from each population at the end of the evolution experiment. All populations had segmental amplifications encompassing *proBA*\* and *rpoS.* The amplicons ranged in size from 9-125 kb and in copy number from 2-12 copies (Fig. 4A and B). Copy numbers were confirmed by quantitative PCR (qPCR) using primers targeting *proA*\*. Most of the segmental amplifications were mediated by the transposable insertion sequence elements IS186 and IS1, leading to similar amplification junctions across populations. IS186 transposition into *serV* served as an upstream junction point in 7 of the 8 populations and IS186 transposition into the intergenic region between *ygcE* and *queE* served as a downstream junction in 6 of the 8 populations (Fig. 4A and *SI Appendix,* Table S1). IS1 transposition was detected at amplification junctions in populations 3 and 7 (*SI Appendix,* Table S1). As neither IS element was present at these sites in the ancestral strain, it appears that these sites may be hotspots for IS element insertion.

**Fig. 4.**
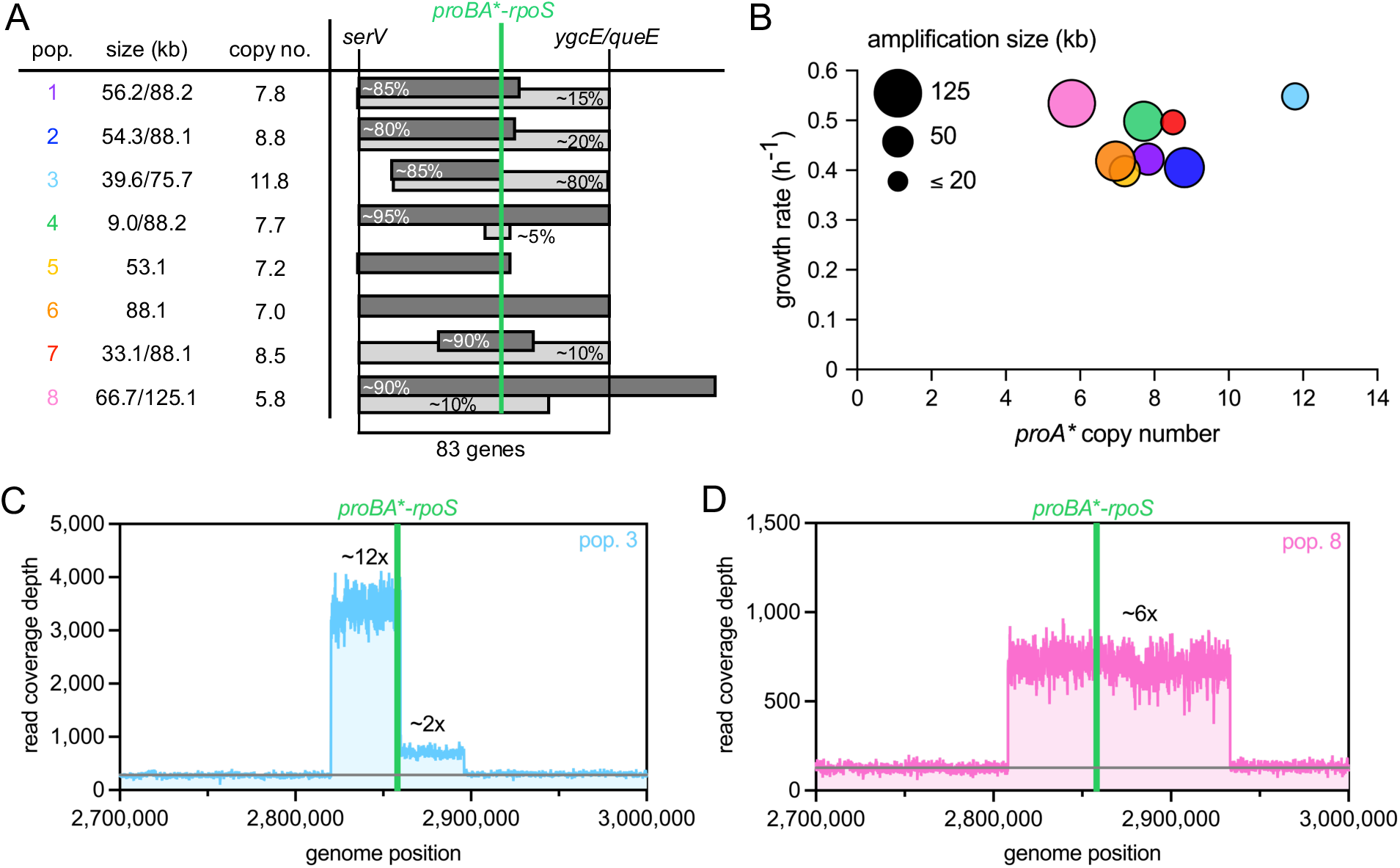
Segmental amplifications occurred during evolution of all *proBA*\*-*rpoS* relocation populations. (A) Sizes and copy numbers of amplicons detected in the eight populations. In populations with two amplicons, dark gray bars represent the higher-frequency amplicon and light gray bars represent the lower-frequency amplicon. Only amplicons estimated to be present at population frequencies of ≥5% are shown. Population-level amplicon frequencies were estimated based on diagnostic PCR for populations 2 and 3 and read coverage data for the remaining six populations. (B) Growth rate, *proA*\* copy number, and amplification size for each population at the end of the evolution experiment. Only the most abundant amplicon is shown for each population for simplicity. (C) The nested amplicons in population 3. (D) The 125 kb amplicon in population 8.

We detected two amplicons at frequencies ≥5% in 6 of the 8 populations based on the genome sequencing data (Fig. 4A). The presence of two amplicons could be due to two competing clades with distinct amplicons in the population or to an amplicon nested inside a larger amplicon in a single clade. Populations 2 and 3 stood out as being the most likely to contain a nested amplicon due to the detection of multiple (≥3) amplification junctions and the presence of amplified regions with distinct copy numbers based on the read coverage data (*SI Appendix,* Fig. S1A and S2A). Thus, we sought to experimentally discern whether there were nested amplicons or competing clades with different amplicons in populations 2 and 3. To distinguish between these possibilities, we carried out PCR reactions designed to amplify products that span putative amplification junctions (*SI Appendix,* Fig. S1 and S2). Population 2 appears to have two distinct clades; almost all colonies (19 out of 20) had either the ∼54 kb (4/19) or the ∼88 kb amplicon (15/19) (*SI Appendix,* Fig. S1). Only one colony (1/20) had both amplicons, suggesting that if a clade with a nested amplicon exists in population 2, it is present at a low frequency. In contrast, the majority of colonies isolated from population 3 (13 out of 20) showed PCR products for both the ∼40 and ∼76 kb amplicons, indicating that most cells contained a smaller approximately 12-copy amplification nested inside a larger duplication (Fig. 4C and *SI Appendix,* Fig. S2). Of the remaining isolates tested from population 3, four had only the ∼40 kb amplicon, while three had only the ∼76 kb amplicon.

We did not perform diagnostic PCRs for the remaining populations with clear high-frequency and low-frequency amplicons based on the read coverage data. In populations 1 and 7, we suspect that low-frequency clades with ∼88 kb amplicons are being outcompeted by clades with smaller amplicons (∼56 and ∼33 kb). In populations 4 and 8, low-frequency clades with ∼9 and ∼67 kb amplicons, respectively, might have arisen more recently than the clades with larger amplicons (∼88 and 125 kb) and thus might still represent a small fraction of the population. Alternatively, the ∼9 and ∼67 kb amplicons might have arisen by remodeling of the larger amplicons in clades that are now in the process of overtaking the original clades.

We focused on populations 3 and 8 for transcriptomic and proteomic analysis because they possessed the highest-copy-number and largest amplifications, respectively (Fig. 4C and D). Most cells in population 3 appear to have a ∼40 kb amplification present at 12 copies nested inside a ∼76 kb duplication (Fig. 4C), while population 8 primarily consists of a clade with a 6-fold amplification of a 125 kb region, the largest amplicon we observed (Fig. 4D). We detected a putative compensatory mutation that was fixed in population 8 at the end of the evolution experiment, which will be discussed in-depth later. Because our goal was to characterize the immediate effects of segmental amplification on the transcriptome and proteome in the absence of compensatory mutations, we sequenced population 8 samples at earlier timepoints and determined that this mutation was not present in any isolates screened at generation 125 (47, 48).

### Expression of most amplified genes is elevated in the aftermath of segmental amplification

We performed RNA-seq and label-free quantitative proteomics on replicate cultures started from single colonies isolated from population 3 at generation 199 and from population 8 at generation 125. We verified that the population 3 isolate contained the ∼40 kb amplification nested inside a larger ∼76 kb duplication by PCR using primers near the amplification junctions. Transcriptomic and proteomic analyses of this isolate allowed us to characterize how mRNA and protein levels changed at two different amplification levels, 2 and 12 copies, within the same genome. The ancestral *rpoS-proBA*\* relocation strain was used as the reference strain for determining expression fold-changes.

Levels of mRNAs and small regulatory RNAs (sRNAs) expressed from amplified genes generally scale linearly with gene copy number (Fig. 5A and *SI Appendix,* Fig. S3, R^2^ = 0.63). On average, RNA levels increased 2.1 ± 0.1-, 5.2 ± 0.2-, and 10.0 ± 0.6-fold (mean ± SEM) for genes present at 2, 6 and 12 copies, respectively (Fig. 5A). Protein levels increased on average by 2.3 ± 0.1-, 5.0 ± 0.2-, and 10.5 ± 1.3-fold for genes present at 2, 6, and 12 copies, respectively (Fig. 5B). However, protein fold-change values varied widely, especially for 6-and 12-copy genes. Levels of proteins expressed from amplified genes correlate relatively weakly with both gene copy number (R^2^ = 0.47) and mRNA level (R^2^ = 0.35) (*SI Appendix,* Fig. S3). Similar results have been observed in aneuploid yeast (37, 47). To some extent the poor correlation may be because estimation of protein abundance by mass spectrometry is less precise than estimation of RNA levels by RNA-seq. However, in many cases, posttranscriptional and posttranslational regulatory mechanisms likely modulate RNA stability, translation and protein degradation in ways that attenuate the impact of increased gene copy number.

**Fig. 5.**
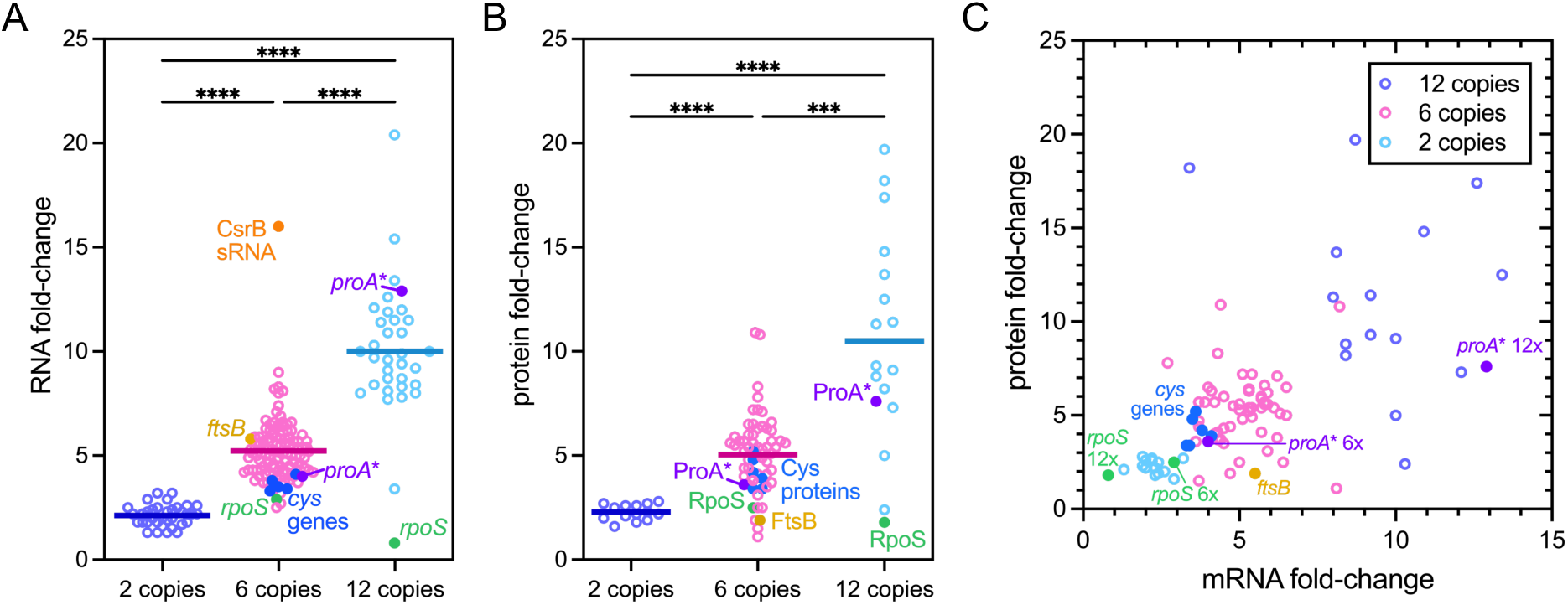
Effect of segmental amplification on expression of RNA and protein from amplified genes. (A) Fold-change of RNAs expressed from amplified genes. (B) Fold-change of proteins expressed from amplified genes. (C) Protein fold-change as a function of mRNA fold-change for amplified genes. Asterisks indicate statistically significant differences between groups based on a non-parametric Kruskal-Wallis test with Dunn’s multiple comparisons test (*** *p* <0.001; **** *p* <0.0001). Amplified genes that are discussed in the main text are indicated by different colors.

### Expression of *rpoS* is attenuated in both populations 3 and 8

The most glaring discrepancy we observed between gene copy number and mRNA and protein levels is the minimal change in *rpoS* mRNA and RpoS protein when *rpoS* is present at 12 copies in population 3 (Fig. 5A and B). The junction between the smaller amplicon and the larger duplicated region in population 3 (Fig. 4C) lies between *rpoS* and *nlpD*; the nested amplification separates *rpoS* from six of seven potential promoters by moving them hundreds of kb upstream of the *rpoS* start codon (Fig. 6A). Thus, population 3 avoids problematic overexpression of *rpoS* even when the gene copy number is very high. Expression of *rpoS* is also not proportional to the 6 copies of the gene in population 8 (Fig. 5C). RpoS levels are tightly regulated by a combination of multiple transcriptional regulators, sRNAs, and an adaptor-mediated proteolysis pathway (45). The combined activity of these regulatory mechanisms may keep RpoS levels in check in the absence of the canonical stresses to which it normally responds.

**Fig. 6.**
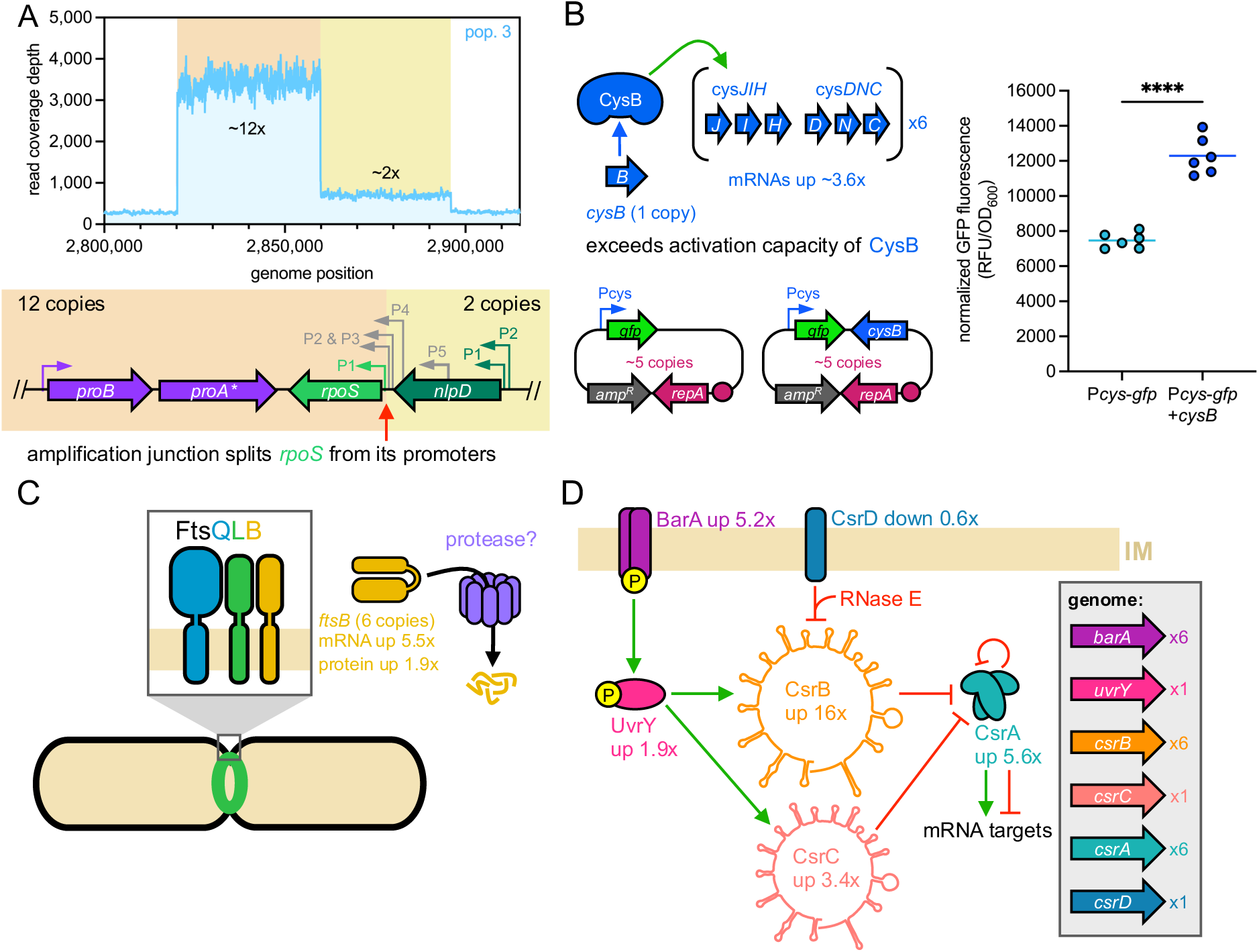
Mechanisms that lead to discrepancies between gene copy number and transcript or protein levels. (A) The nested amplification junction present in population 3 separates *rpoS* from 6 of 7 potential promoters. (B) Normalized GFP fluorescence for cells carrying a plasmid with a copy number of 5 encoding *gfp* under control of a CysB-activated promoter with and without *cysB* on the same plasmid. Asterisks indicate statistically significant differences between groups based on an unpaired *t* test (**** *p* <0.0001); n=6 biological replicates. (C) Lower-than-expected FtsB protein levels result from decreased FtsB stability and thus more degradation in the absence of enough of its interaction partners FtsQ and FtsL. (D) Disproportionate expression of genes involved in the Csr system results from both amplification of some but not all genes and complex interactions between RNA and protein components of the circuit.

### Amplification of *cysJIH-DNC* outstrips the availability of the transcriptional activator CysB

The *cysJIH* and *cysDNC* operons, which encode proteins involved in sulfate assimilation, were amplified to 6 copies in population 8, but levels of the encoded mRNAs increased by an average of only 3.6-fold (Fig. 5C and Dataset S1). Expression of these operons requires the transcriptional activator CysB, which is encoded outside the amplified region and is thus present at only a single copy (Fig. 6B). We hypothesized that the relatively modest transcriptional response of the *cysJIH* and *cysDNC* operons following amplification was due to insufficient CysB to initiate transcription at all of the extra copies of the *cysDNC* and *cysJIH* operons. To test this hypothesis, we constructed a plasmid-based transcriptional reporter system with GFP expression under the control of a CysB-activated promoter (the promoter of the *cysDNC* operon) (Fig. 6B). We constructed two versions of this low-copy-number reporter plasmid (∼5 copies per cell), one lacking *cysB* and another encoding *cysB.* Cells harboring the reporter plasmid lacking *cysB* rely on the single chromosomal copy of *cysB* to activate *gfp* transcription. Conversely, cells harboring the plasmid containing *cysB* have the chromosomal copy of *cysB* along with ∼5 extra copies of *cysB* from the plasmid, a scenario in which the copy number of *cysB* is commensurate with the copy number of the *cysDNC* and *cysJIH* operons. We found that cells with the reporter plasmid containing *cysB* had 1.6-fold higher GFP fluorescence than cells lacking *cysB* on the reporter plasmid (Fig. 6B). This finding suggests that the dampened *cysDNC* and *cysJIH* transcript levels after amplification are due to outstripping of the activation capacity of CysB.

### Dosage imbalance leads to degradation of a protein produced at a higher level than its interaction partners

FtsB is an essential cell division protein that forms a 1:1:1 complex with FtsL and FtsQ (48). *ftsB* is located within the 6-fold amplified region in population 8, but *ftsL* and *ftsQ* are over 2.7 Mb away from *ftsB* and not amplified. *ftsB* expression was increased by 5.5-fold at the transcript level, but only 1.9-fold at the protein level (Fig. 5C and Dataset S1). Proteins that form complexes are often unstable in the absence of their interaction partners due to improper folding and subsequent degradation by proteases (49, 50). It has been shown that FtsL and FtsQ can both stabilize FtsB, preventing its degradation (51). Depletion of either FtsL or FtsQ in *E. coli* resulted in reduced levels of FtsB and simultaneous appearance of a smaller FtsB degradation product (51). Thus, the imbalance between the cellular levels of FtsB and FtsLQ we observe after co-amplification of *ftsB* likely leads to decreased stability and increased degradation of FtsB protein without affecting *ftsB* transcript levels, which scale with gene copy number (Fig. 6C).

### The structure of the Csr circuit seems to buffer it against dosage imbalances caused by segmental amplification

In population 8, *csrA*, *csrB* and *barA* are present within the 6-copy amplicon. The levels of CsrA and BarA proteins are increased by 5.6- and 5.2-fold, respectively. However, the level of the sRNA CsrB is disproportionally increased by 16-fold (Fig. 6D and Dataset S1). These genes are components of the carbon storage regulator (Csr) system, a complex regulatory circuit consisting of both proteins and sRNAs that posttranscriptionally coordinates expression of genes involved in diverse processes such as glycolysis, glycogen synthesis and flagellar motility (52, 53). *csrC*, *csrD* and *uvrY*, which are also part of the Csr system, were not amplified, but the levels of the sRNA CsrC and the protein UvrY are increased by 3.4- and 1.9-fold respectively, while the level of CsrD is modestly decreased by ∼40% (Fig. 6D and Datasets S2 and S3). Thus, amplification of some Csr system genes appears to lead to perturbations in the levels other components.

The striking and concurrent increases in the levels of CsrB and CsrC sRNAs in population 8 are likely due to a combination of effects mediated by the intricate network of regulatory interactions in the Csr circuit (Fig. 6D). CsrB and CsrC are functionally similar, as both sRNAs can bind and sequester CsrA homodimers in an inactive state (52, 53). Despite *csrB* being within the amplified region, while *csrC* is located >1.1 Mb away and not amplified, both sRNAs are largely subject to the same forms of regulation in the Csr system. The 5.2-fold increase in the level of the histidine kinase BarA and the 1.9-fold increase in the level of the cognate response regulation UvrY would be expected to increase transcription of *csrB* and *csrC.* In addition, the decreased expression of CsrD, which recruits RNase E to degrade CsrB and CsrC sRNAs, should also contribute to increased levels of CsrB and CsrC.

To our surprise, the substantial perturbations to sRNA and protein levels in this elaborate regulatory circuit seem to have negligible effects on the stability and translation of mRNAs regulated by CsrA. CsrA blocks translation of *glgCAP* mRNAs, which encode glycogen synthesis enzymes (52, 53). However, we see little change in GlgCAP protein levels (Dataset S2). Conversely, CsrA protects *flhDC* mRNAs, which encode the master regulator of flagellar synthesis, from degradation by RNAse E, thereby promoting translation (52, 53). Although *flhDC* transcript levels are slightly elevated, we did not detect FlhDC proteins in our proteomics data (Dataset S2 and S3). We observed a similar modest increase in *flhDC* transcript levels in population 3, in which neither *csrA* nor *csrB* is amplified, suggesting that CsrA plays little role in altering *flhDC* mRNA levels in these strains. The effect of the elevated CsrA expression is likely blunted by the increased levels of both CsrB and CsrC, which sequester CsrA in an inactive form. Thus, the net result of moderately elevated CsrA levels may be insignificant in the context of other regulators that modulate transcription of *glgCAP* and *flhDC* (54). Overall, the intricate wiring of the Csr regulatory circuit, replete with different forms of activation and inhibition, seems to buffer it against stoichiometric imbalances between interacting sRNAs and proteins. This intrinsic buffering capacity should be useful for acclimating to environmental changes while avoiding dysregulation, but it also seems to have the incidental effect of limiting severe physiological perturbations following segmental amplification.

### Segmental amplification has system-wide effects

The segmental amplifications in populations 3 and 8 resulted in significant changes hundreds of transcripts (Fig. 7), including mRNAs and sRNAs encoded by genes that were not present in the amplified regions. Expression of two classes of non-amplified genes was clearly increased in both populations 3 and 8: ribosomal RNA (rRNA) and ribosomal protein genes; and colanic acid capsular polysaccharide synthesis and export genes (Fig. 7 and Dataset S3).

**Fig. 7.**
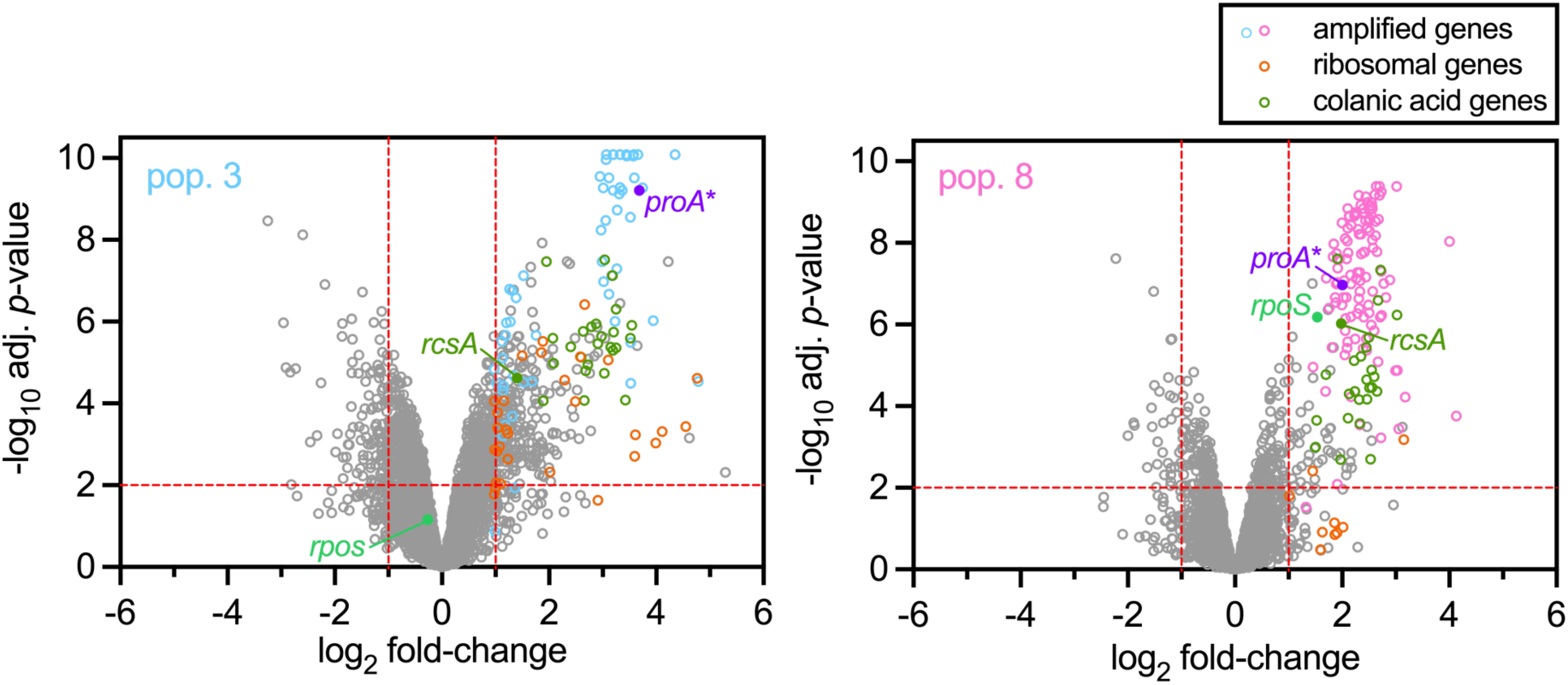
The global transcriptomic consequences of segmental amplification in populations 3 and 8. Amplified genes that are differentially expressed in populations 3 and 8 compared to the ancestral strain are indicated by blue and pink points, respectively. Amplified genes *proA*\* and *rpoS* and classes of non-amplified genes are indicated by different colors.

The increased expression of rRNA and ribosomal protein genes is likely an indirect effect of the amplification of *proA*\*, which alleviates the shortage of proline and arginine in the ancestral strain and should dampen the stringent response that diminishes transcription of ribosomal genes (55). The increased expression of about 25 colanic acid capsular polysaccharide synthesis genes appears to be caused by overexpression of the transcriptional activator RcsA. Although *rcsA* is not among the co-amplified genes, *rcsA* transcript levels are increased by 2.6- and 4-fold and protein levels are increased by 8.2- and 4.6-fold in populations 3 and 8, respectively (Fig. 7 and Datasets S2 and S3). RcsA forms a heterodimer with RscB that stimulates transcription of *rcsA* itself in addition to colanic acid synthesis genes (56–58). RcsA is an unstable protein that is rapidly degraded by Lon and HslUV proteases, which typically prevents excess RcsA from accumulating and activating its own transcription(56–60). Because proteolysis appears to be the primary mechanism by which RcsA expression is kept low, a physiological perturbation that impairs global protein homeostasis might be expected to attenuate the capacity of Lon and HslUV proteases to degrade RcsA. We hypothesize that inappropriate overexpression of co-amplified genes following segmental amplification might divert Lon and other proteases away from degrading their usual client proteins, such as RcsA, and instead shift their activity toward degrading overexpressed and potentially misfolded proteins. The resulting accumulation of RcsA would, in turn, increase transcription of *rcsA* itself as well as colanic acid synthesis genes.

### A compensatory mutation elevates Lon protease activity and alleviates adverse effects of segmental amplification

As mentioned previously, we identified a putative compensatory mutation that was fixed in population 8 at the end of the evolution experiment. This mutation is a small 7 bp duplication in *prlF*, which encodes the antitoxin of the PrlF-YhaV toxin-antitoxin system (61). This exact mutation has been reported to alleviate toxic overexpression of a fusion protein by hyperactivating the Lon protease (61, 62). While the molecular mechanism linking this toxin-antitoxin system to the activity of Lon protease is not clear, this *prlF* mutation may help ameliorate general proteotoxic stress, as Lon plays an important role in global protein homeostasis by degrading misfolded proteins in addition to particular client proteins (63, 64). This *prlF* mutation was not detected in colonies isolated from population 8 at generation 125 yet rose to fixation in approximately 60 generations by generation 186. The highly beneficial nature of this *prlF* mutation is consistent with the notion that increasing protein degradation by elevating Lon activity alleviates the some of the fitness cost associated with segmental amplification.

Our initial transcriptomic and proteomic analysis of population 8 used a clonal isolate from generation 125, which lacked the *prlF* mutation, in order to characterize the immediate collateral consequences of segmental amplification in the absence of compensatory mutations. To determine the global effect of the Lon-hyperactivating *prlF* mutation, we also isolated a clone from population 8 at generation 186 with the *prlF* mutation and performed RNA-seq and label-free quantitative proteomics in parallel with our other analyses. The population 8 isolates from generations 125 and 186 have the same ∼125 kb amplicon at approximately 6 copies (*SI Appendix* Fig. S4), and the only difference between the two isolates is the *prlF* mutation. The *prlF* mutation causes a significant decrease in the mRNA levels of many colanic acid synthesis and export genes, including *rcsA* (Fig. 8A). Because the *prlF* mutation increases the activity of Lon protease and Lon is known to readily degrade RcsA (56–58), lower RcsA levels are expected to result in less transcription of colanic acid-related genes.

**Fig. 8.**
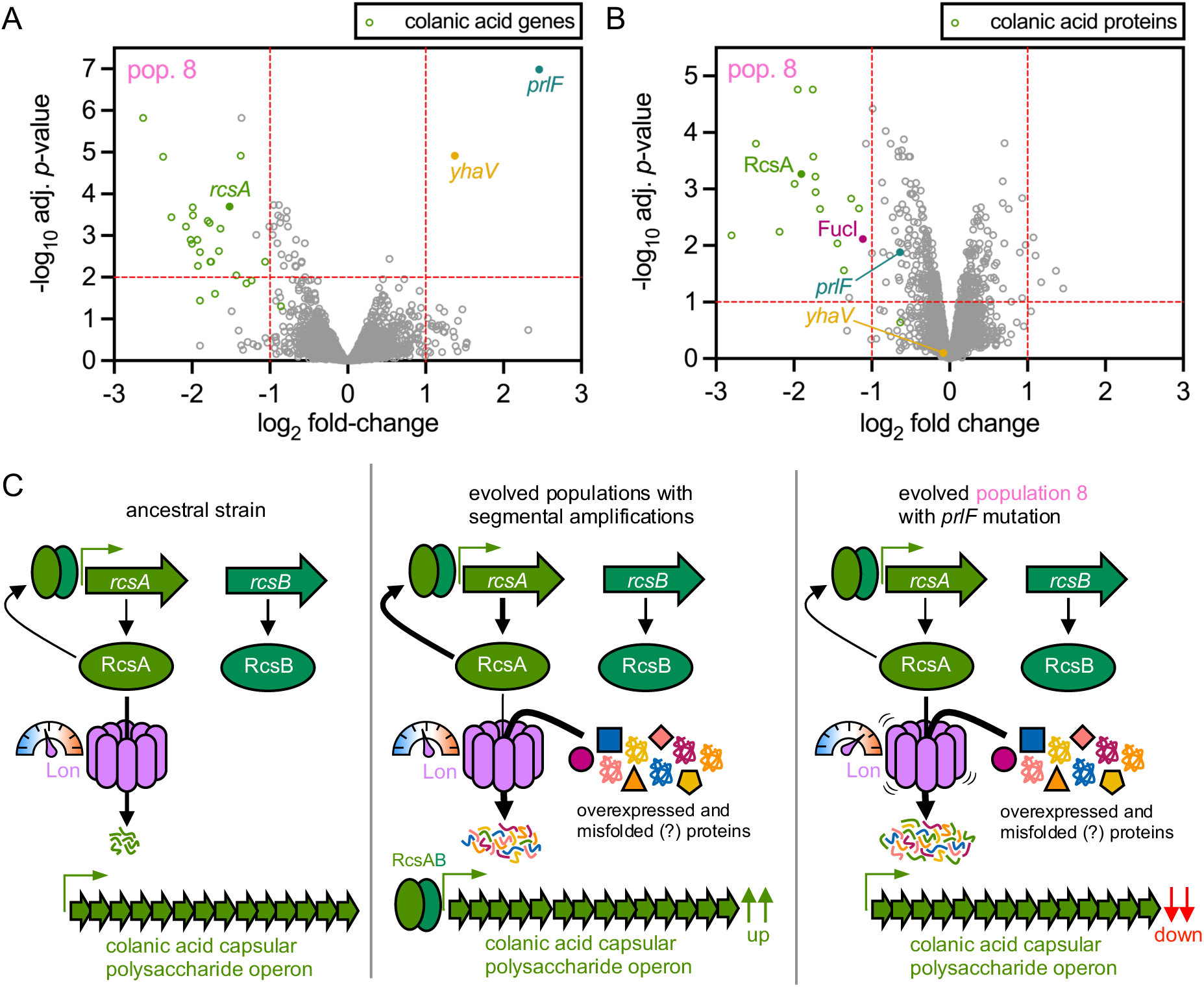
A compensatory mutation that increases activity of Lon protease decreases expression of colonic acid synthesis and export genes. (A) The effect of the *prlF* mutation on the transcriptome of population 8. (B) The effect of the *prlF* mutation on the proteome of population 8. Gene expression was compared between a generation 125 isolate lacking the *prlF* mutation and a generation 185 isolate with the *prlF* mutation. Colanic acid gene/proteins are indicated in green. Other genes/proteins discussed in the text are indicated with different colors. (C) Postulated effect of the compensatory *prlF* mutation that hyperactivates Lon protease, leading to increased degradation of RcsA and therefore decreased expression of RcsAB-activated colanic acid capsular polysaccharide biosynthesis genes.

Only two genes, *prlF* itself and *yhaV*, which together form an operon and are thus transcribed as a single polycistronic mRNA (61), show dramatic upregulation due to the *prlF* mutation. The PrlF antitoxin and YhaV toxin together form a transcriptional repressor that represses transcription of their own operon (61). The *prlF* mutation is a 7 bp duplication that causes a frameshift, which truncates the C-terminus of PrlF and impairs the repressor function of PrlF-YhaV (61). Thus, diminished repression of the *prlF-yhaV* operon due to the *prlF* mutation increases transcript levels for these genes.

At the proteome level, we observe a clear decrease in the expression of colanic acid synthesis and export proteins, matching their dampened expression at the transcript level (Fig. 8A and B). Our data support the hypothesis that the *prlF* mutation causes a massive attenuation of many colanic acid-related genes at the transcript and protein levels by increasing Lon-mediated proteolysis of the transcriptional activator RcsA. Increasing degradation of RcsA alone likely prevents the unnecessary expression of over 20 genes involved in colanic acid synthesis and export. We also wondered whether increasing Lon activity via the *prlF* mutation might decrease the inappropriate overexpression of any co-amplified genes. The level of FucI, a fucose isomerase, is increased by 7.2-fold in the generation 125 isolate with 6 copies of *fucI* compared to the ancestral strain (Dataset S1). In the generation 186 isolate with the *prlF* mutation, FucI levels are reduced by >50% despite *fucI* still being present at 6 copies (Fig. 8B). Because fucose is not present in the growth medium, FucI expression is not needed for growth. The reduction in FucI levels suggests that the hyperactivated Lon in the generation 186 isolate contributes to FucI degradation. However, other proteins encoded in the amplified *fucAO* and *fucPIKUR* operons do not exhibit similar decreases in expression following hyperactivation of Lon (Dataset S1).

While the transcription of the *prlF-yhaV* operon is highly upregulated due to the *prlF* mutation, their expression at the protein level is not (Fig. 8 A and B). This result is not surprising as the 7 bp duplication in *prlF* introduces a premature stop codon that has been shown to have polar effects on *yhaV*, possibly by decreasing translation of YhaV from polycistronic *prlF-yhaV* mRNAs (61). The truncated PrlF that results from this frameshift mutation has a shorter and more basic C-terminus. The altered C-terminus of PrlF and/or diminished interaction with YhaV may reduce PrlF stability and therefore its level (Fig. 8B).

The effect of the *prlF* mutation on the transcriptome and proteome of cells with large segmental amplifications indicates that it is a compensatory mutation. We hypothesize that by elevating the activity of Lon protease, the *prlF* mutation helps alleviate the fitness costs associated with inadvertent overexpression of amplified and non-amplified genes (Fig. 8C). In population 8, the *prlF* mutation, which increases Lon protease activity (62), arises and fixes in about 60 generations, indicating it is highly beneficial. When comparing isolates with and without the *prlF* mutation, our data show that hyperactivating Lon leads to more degradation of client protein RcsA and thus less transcription and lower protein levels of colanic acid synthesis genes (Fig. 8). Therefore, we view the *prlF* mutation as representing a class of compensatory mutations that are broadly beneficial for counteracting proteotoxic stress by increasing proteolysis, which could result from segmental amplification or other forms of stress that disrupt protein homeostasis.

## Summary

The strong selection for increased ProA* activity in our evolving populations resulted in rapid segmental amplification encompassing *proA*\* and many neighboring genes. Transcription of most amplified genes correlates well with copy number. However, there are notable exceptions. Expression of *rpoS* is attenuated in population 3 by loss of several of its promoters. Excessive RpoS would be expected to be problematic because RpoS promotes a transcriptional program that slows growth and alters resource allocation when cells are starving or stressed. This response would be inappropriate when glucose is abundant and proline and arginine synthesis have been improved due to amplification of *proA*.* It is tempting to speculate that the loss of *rpoS* promoters allowed the small amplicon nested within the originally duplicated region to rise to 12 copies.

We also observed other mechanisms for attenuation of mRNA expression that do not rely on a fortuitous amplification junction. Transcription of amplified genes in two operons encoding sulfate assimilation genes is attenuated due to inadequate levels of a transcriptional activator encoded by a gene that was not amplified. Such a mismatch could occur in many situations when only some genes in a regulon are amplified. Regulons often include operons that are scattered across the genome and are controlled by transcriptional activators that may not be in close proximity to the operons they control. While attenuation in this case appears to be due to outstripping the capacity of a transcriptional activator, we expect that the opposite situation could also arise; i.e. expression of amplified genes could be exacerbated if the capacity of a transcriptional repressor is outstripped.

Post-transcriptional mechanisms also appear to contribute to attenuation of protein levels produced from amplified genes. For example, it is well accepted that subunits of protein complexes that are produced in excess are often degraded, probably because they misfold due to exposure of surfaces that are normally involved in protein-protein interactions (49, 50). Indeed, excess FtsB has been shown to be degraded due to insufficient levels of its interacting partner proteins FtsQ and FtsL (51).

The complex regulation of many genes in *E. coli* may contribute to attenuation of expression of amplified genes. *rpoS* expression is regulated by multiple transcription factors, sRNAs, and adaptor-mediated proteolysis, which allows integration of multiple signals in the decision to activate the stress response while preventing inopportune overexpression. Likewise, the Csr system integrates multiple inputs in a complex regulatory circuit in order to appropriately fine-tune translation of specific mRNAs. While these systems evolved to enable *E. coli* to respond appropriately to changing environments, a propitious side effect is the ability to buffer the effects of segmental amplification.

Segmental amplification can have global consequences. Approximately 86% and 49% of genes whose expression is increased by ≥2-fold at the transcript level in populations 3 and 8, respectively, are not amplified. While co-amplification of genes neighboring a gene under selection for increased copy number has previously been recognized to contribute to fitness, our work adds a previously unappreciated nuance; segmental amplification can also alter expression of mRNAs and proteins from hundreds of unamplified genes. Thus, the immediate fitness effects of segmental amplification are due to the combination of the desirable effect of amplifying the gene under selection for increase dosage and the collateral effects (both detrimental and beneficial) of altered expression of co-amplified genes within the amplicon as well as genes elsewhere in the genome. Finally, the fitness costs associated with segmental amplification can be mitigated, at least partially, by compensatory mutations that alleviate the energetic burden or physiological perturbations that accompany a particular amplification.

## Materials and Methods

### Strains and growth conditions

*E. coli* strains, plasmids, and primers used in this work are listed in *SI Appendix*, Tables S2, S3, and S4. *E. coli* strains were routinely cultured in lysogeny broth (LB) or on LB agar plates containing 1.5% agar at 37 °C. M9 salts minimal medium (12.8 g/L Na_2_HPO_2_•7H_2_O, 3 g/L KH_2_PO_4_, 0.5 g/L NaCl, 1 g/L NH_4_Cl) was supplemented with glucose at a final concentration of 0.2% (v/v). Antibiotics were used at the following final concentrations unless otherwise indicated: ampicillin, 100-150 μg/mL; kanamycin, 20 μg/mL; chloroamphenicol, 20 μg/mL; and gentamicin, 15 μg/mL.

### Strain construction

All strains used in this work are derived from *E. coli* BW25113. The Δ*argC*::*kan^R^* M2-*proBA*\*-*yfp* Δ*fimAICDFGH* Δ*csgBAC* Δ82 bp *rph-pyrE E. coli* strain AM327 was previously constructed from the Keio collection Δ*argC*::*kan^R^ E. coli* BW25113 strain as described in (4). The Δ*fimAICDFGH* Δ*csgBAC* deletions help prevent biofilm formation during laboratory evolution in a turbidostat. The 82 bp deletion in *rph* and upstream of *pyrE* fixes a defect in pyrimidine synthesis due to a frameshift in *rph* in *E. coli* BW25113 and other K-12 strains (65, 66). The M2 promotor mutation is a C > T point mutation 45 bp upstream of *proB* that increases transcription of the *proBA* operon as described in (44). The AM187 strain (GenBank accession number CP037857.2) and its derivative AM327 contain a 1.8-Mb inversion in the genome that occurred between two IS3 elements (43).

For this study, we deleted the M2-*proBA*\*-*yfp* region in the *E. coli* AM327 genome using λ-Red recombineering (67). We transformed the temperature-sensitive plasmid pSIM6 (68) containing the λ-Red recombinase genes (*exo*, *beta* and *gam*) under control of a heat-inducible promoter into AM327 by electroporation. After recovery in super optimal broth with catabolite repression medium (SOC) (BD Difco™, Franklin Lakes, NJ) for ∼3 h at 30 °C, cells were plated on LB ampicillin and incubated at 30 °C to select for transformants. We replaced *proBA\** and *yfp* in AM327 with a *sacB-gtm^R^*selection-counterselection cassette amplified from the plasmid pJQ200SK (69). We amplified ∼1 kb fragments from the regions upstream and downstream of the *proBA* operon using primers that introduced ∼20 bp ends complementary to the ends of the *sacB-gtm^R^* cassette. We assembled a linear *proBA*-yfp* replacement fragment from these three fragments using splicing by overlap extension (SOE) PCR using Q5^®^ High-Fidelity DNA Polymerase (New England Biolabs, Ipswich, MA). The AM327 strain harboring pSIM6 was inoculated into LB ampicillin, grown to an optical density at 600 nm (OD_600_) of 0.2-0.6 at 30 °C and then incubated at 42 °C for 15 min to induce expression of the λ Red genes. Cells were then centrifuged at 4,500 x g at 4 °C, washed with ice-cold deionized water and immediately electroporated with ∼0.5-2.0 μg of the *proBA\** replacement linear fragment. Following electroporation, cells were inoculated into fresh SOC (BD Difco™) and allowed to recover at 30 °C for ∼3 h before being spread onto LB gentamicin. Gentamicin-resistant colonies were screened for integration of *sacB-gtm^R^* into the *proBA* locus by colony PCR using One*Taq*^®^ DNA Polymerase (New England Biolabs).

We constructed a fragment to delete the *sacB-gtm^R^* cassette by amplifying ∼1 kb regions upstream and downstream of the *proBA* operon with overlapping ends. The two PCR products were assembled together using SOE PCR. Cells containing pSIM6 and the *sacB-gtm^R^* selection-counterselection marker in place of *proBA\** were inoculated into LB ampicillin, grown to an OD_600_ of 0.2-0.6 at 30 °C and subjected to a second round of λ-Red recombineering as described above. Cells were electroporated with ∼0.5-2.0 μg of the *sacB-gtm^R^* replacement fragment. Following electroporation, cells were inoculated into fresh SOC and allowed to recover at 30 °C for ∼3 h before being spread onto salt-free (i.e. no NaCl) LB plates containing 10% (w/v) sucrose and incubated at 30 °C for counterselection. Colonies that grew on salt-free LB sucrose plates were patched onto LB gentamicin plates to confirm gentamicin sensitivity due to loss of the *sacB-gtm^R^* cassette and screened for markerless deletion of *proBA\** by colony PCR and Sanger sequencing.

To relocate the M2-*proBA*\* operon next to *rpoS* in the *E. coli* genome, we first introduced a *tetR-ccdB-cat* selection-counterselection cassette at the target site between *rpoS* and *ygbN.* We amplified the *tetR-ccdB-cat* cassette from pDLM3 (70). We amplified ∼1 kb fragments upstream and downstream of the target site using primers that introduced ∼20 bp ends complementary to the ends of the *tetR-ccdB-cat* cassette. These three fragments were assembled together using SOE PCR. The Δ*proBA*\* strain RF72 harboring pSIM6 was inoculated into LB ampicillin, grown to an OD_600_ of 0.2-0.6 at 30 °C and subjected to λ-Red recombineering as described above. Cells were electroporated with ∼0.5-2.0 μg of the *tetR-ccdB-cat* integration fragment. Following electroporation, cells were inoculated into fresh SOC and allowed to recover at 30 °C for ∼3 h or at room temperature overnight before being spread onto LB chloramphenicol plates. The plates were incubated at 30 °C for 1-2 days. Chloramphenicol-resistant colonies were screened for integration of *tetR-ccdB-cat* between *ygbN* and *rpoS* by colony PCR. The *tetR-ccdB-cat* cassette was then replaced with M2-*proBA*.* We amplified the M2-*proBA*\* operon (without *yfp*) from AM187 gDNA and ∼1 kb fragments upstream and downstream of the intergenic site between *rpoS* and *ygbN* from RF72 gDNA by PCR. These three fragments were assembled using SOE-PCR to generate the *tetR-ccdB-cat* replacement cassette. The *ygbN-tetR-ccdB-cat-rpoS* strain harboring pSIM6 was inoculated into LB ampicillin, grown to an OD_600_ of 0.2-0.6 at 30 °C and subjected to λ-Red recombineering as described above. Cells were electroporated with ∼0.5-2.0 μg of the *tetR-ccdB-cat* replacement cassette. Following electroporation, cells were inoculated into fresh SOC and allowed to recover at 30 °C for ∼3 h or at room temperature overnight before being spread onto LB anhydrotetracycline (2-6 ug/mL) plates for counterselection. The plates were incubated at 30 °C overnight. Colonies that grew on LB anhydrotetracycline plates were patched onto LB chloramphenicol plates to confirm chloramphenicol sensitivity and screened for integration of M2-*proBA*\* into the intergenic site between *ygbN* and *rpoS.* The resulting M2-*proBA*\*-*rpoS* strain was streaked onto LB anhydrotetracycline plates and incubated at 40 °C overnight to cure the temperature-sensitive pSIM6 plasmid. Colonies were patched onto fresh LB anhydrotetracycline and LB ampicillin plates to confirm ampicillin sensitivity due to loss of pSIM6. Genomic integration of M2-*proBA*\* next to *rpoS* was confirmed by Sanger sequencing.

### Laboratory Evolution

To initiate laboratory evolution of the *proBA*\*-*rpoS* relocation strain (RF90), cells from a frozen glycerol stock were streaked onto LB kanamycin plates and the plates were incubated at 37 °C overnight. A single founder colony was inoculated into LB kanamycin and incubated at 37 °C overnight. The overnight culture was centrifuged at 8,000 x g at room temperature and the cells were washed and resuspended in M9 medium without glucose. A 125 μL aliquot of cells was inoculated into each of eight glass vials containing 25 mL (1:200 dilution) of M9 minimal medium containing 0.2% glucose and kanamycin (20 µg/mL). The vials were placed into the vial sleeves of an eVOLVER turbidostat (FynchBio). Each vial contained a micro-flea magnetic stir bar (10 mm x 3 mm). The eVOLVER was set in ‘turbidostat’ operation mode at 37 °C with a stir rate of 8, a lower OD_600_ threshold of 0.15, an upper OD_600_ threshold of 0.25 and a vial volume of 25 mL. When cultures reached the upper OD_600_ threshold of 0.25, ∼7-10 mL of fresh M9 + glucose medium was pumped into vials and an equivalent volume of culture was removed so that cultures were diluted to an OD_600_ of ∼0.15. Cultures were evolved in this manner for 145-200 generations (∼14 days). Approximately every 40-60 h, ∼7-10 mL samples of cultures expelled from the vials during dilution were collected and centrifuged at 4,500 x g at room temperature. The cell pellets were resuspended in 1 mL of M9 medium without glucose. Aliquots (0.5 mL) of concentrated cells from each population were mixed 1:1 with 50% glycerol and stored at -70 °C. The remaining 0.5 mL cell suspensions were centrifuged at 8,000 x g at room temperature, the supernatants were removed and the cell pellets were stored at -20 °C for extraction of genomic DNA (gDNA).

### Whole Genome Sequencing and Analysis

Cell pellets were thawed and gDNA was extracted using the PureLink™ Genomic DNA Mini Kit (Invitrogen) according to the manufacturer’s protocol for Gram-negative bacteria. Purified gDNA samples were stored at -20 °C. Whole genome sequencing was performed at SeqCenter (formerly the Microbial Genome Sequencing Center; Pittsburgh, PA). Sample libraries were prepared with the Illumina DNA Prep kit and IDT 10 bp UDI indices and sequenced on an Illumina NextSeq 2000, producing 2×151 bp paired-end reads. Demultiplexing, quality control, and adapter trimming was performed with Illumina BCL Convert v3.9.1. The resulting fastq files were further processed using fastp v0.20.1 (71) to trim and remove low quality reads with the following settings: -q 15 -u 20 -t 1. Mutations in evolved populations were identified using breseq v0.37.1 (72, 73) in polymorphism mode with bowtie 2.5.0 (74) and R 4.2.2 by mapping processed fastq reads to the genome of the *E. coli proBA*\*-*rpoS* strain. To visualize read coverage depth of segmental amplifications encompassing *proBA*\*, read coverage depth for every 50 bp for a 300 kb genome window was exported using the breseq BAM2COV subcommand: -a seq id:2700000-3000000 -t -1 --resolution 6000.

### Quantitative PCR for measurement of *proA*\* copy number

*proA*\* copy number was measured using qPCR of purified gDNA as described in (4) with slight modifications. Briefly, PowerSYBR Green PCR master mix (Applied Biosytems by Thermo Scientific) was combined with primers and gDNA in 48-well plates (Thermo Scientific) and samples were analyzed using a StepOne Real-Time PCR System (Applied Biosystems). The genes *gyrB* and *icd*, which remained at one copy in evolved populations, were used as internal reference genes for quantification of *proA*\* copy number. Threshold cycle (C_T_) measurements for duplicate technical replicates of each gDNA sample were averaged. The ΔC_T_ values for *proA*\* and single copy reference genes were calculated by subtracting the C_T_ for gDNA from the ancestral *proBA*\*-*rpoS* strain from the C_T_ for gDNA from the evolved populations.

The ΔC_T_ values of reference genes *gyrB* and *icd* were averaged for each sample and ΔΔC_T_ values were then calculated by subtracting the mean reference ΔC_T_ from *proA*\* ΔC_T_ for each sample. *proA*\* copy number was calculated as 2^-ΔΔCT^ for each sample. Primer sets used for qPCR are listed in *SI Appendix* Table S4.

### RNA extraction

Glycerol stocks of ancestral *proBA*\*-*rpoS E. coli* and evolved populations 3 (at generation ∼199) and 8 (at generation ∼125) were streaked onto M9 plates containing 0.4% glucose. Single colonies of each strain were inoculated into 4 mL M9 containing 0.2% glucose and the cultures were grown at 37 °C overnight with shaking at 200 rpm. The overnight cultures were inoculated into 25 mL of fresh M9 containing 0.2% glucose and 20 μg/mL kanamycin in 125 mL Erlenmeyer flasks at a 1:50 dilution for the ancestral strain and a 1:100 dilution for the evolved populations to account for differences in growth rate. Four replicate cultures of each strain were grown at 37 °C with shaking at 200 rpm for ∼7 h to an OD_600_ of ∼0.2-0.5. Cultures were centrifuged at 4,500 x g for 10 min at 4 °C. Cell pellets were resuspended in 5 mL M9 salts without glucose. Four mL of each cell suspension was flash frozen in a dry ice-ethanol (95%) bath and stored at -70 °C prior to RNA extraction. The remaining 1 mL of each cell suspension was centrifuged at 8,000 x g for 2 min. The supernatant was removed and the cell pellets were flash frozen in a dry ice-ethanol (95%) bath and stored at -20 °C prior to sample preparation for proteomics.

Total RNA was extracted from cells using RNAprotect Bacteria Reagent and RNeasy Mini Kit (Qiagen, Hilden, Germany). Frozen 4-mL cell suspensions were thawed on ice, mixed 2:1 with RNAprotect Bacteria Reagent, incubated for 5 min at room temperature and centrifuged at 4,500 x g for 10 min at room temperature. Cell pellets were resuspended in TE buffer with lysozyme (1 mg/mL) to lyse the cells. RNA was purified using RNeasy Mini Kits according to the manufacturer’s instructions except that optional on-column DNase digestion was performed by mixing 10 μL TURBO™ DNase (Invitrogen, Waltham, MA) with 70 μL buffer RDD (Qiagen), adding 80 μL of DNase solution directly to the spin column membrane of each sample and incubating at room temperature for 15 min. RNA samples were stored at -20 °C.

### RNA sequencing and differential expression analysis

RNA sequencing was performed at SeqCenter (formerly the Microbial Genome Sequencing Center; Pittsburgh, PA). Three replicate RNA samples were sequenced for each strain. RNA samples were treated with DNase (Invitrogen). Sample libraries were prepared with the Illumina Stranded Total RNA Prep Ligation with Ribo-Zerp Plus kit and 10 bp IDT for Illumina indices and sequenced on a NextSeq 2000, producing 2×51 bp paired-end reads. Demultiplexing, quality control, and adapter trimming was performed with Illumina BCL Convert v3.9.1.

To map reads to the *E. coli* genome to generate read counts for each gene, fastq files were processed with fastp v0.20.1 (71) to trim and remove low quality reads with the following settings: -q 15 -u 20. Processed reads were aligned to the *E. coli* reference genome using bowtie2 (74). Read alignment files were converted into sorted.bam files using samtools (75, 76). Aligned reads were mapped to *E. coli* genes using HTSeq (77) to generate read counts with the settings: htseq-count -f bam –r pos -s no -t gene -i Name -m intersection-nonempty --nonunique all.

Differential expression analysis was performed in R v4.3.1/RStudio (Posit team; Boston, MA) with the following Bioconductor/R packages: edgeR (78, 79), limma (80), glimma, gplots, RColorBrewer and NFM. Read counts generated from HTSeq were filtered to remove lowly expressed genes and normalized using trimmed mean of M values with edgeR. Read counts were transformed into log counts per million (CPM) with the voom function of limma for each sample. Differential expression analysis of transformed logCPM data was performed with the contrasts.fit and eBayes functions in limma. Differential expression data were exported and plotted in GraphPad Prism v10.0.2 (GraphPad Software; Boston, MA)

### Label-free proteomics and differential expression analysis

Frozen cell pellets were lysed in 50 mM Tris-HCl, pH 8.5 containing 5% (w/v) sodium dodecyl sulfate, 10 mM tris(2-carboxyethylphosphine) and 40 mM chloroacetamide by boiling for 10 minutes at 95°C followed by probe sonication (1 sec on, 1 sec off) for 1 minute. Each sample was digested using the SP3 method described in (81). Carboxylate-functionalized SpeedBeads (GE Life Sciences) were added to the lysates. Addition of acetonitrile to 80% (v/v) caused proteins to bind to the beads. The beads were washed twice with 80% (v/v) ethanol and twice with 100% acetonitrile. Proteins were digested with Lys-C/Trypsin (Promega) at 37 °C overnight with rotation. SpeedBeads were collected by centrifugation and placed on a magnet to hold the SpeedBead pellet together so that the digested peptides in the supernatant could be extracted. The peptides were then desalted using an Oasis HLB cartridge (Waters) according to the manufacturer’s instructions and dried in a SpeedVac.

The dried peptide samples were suspended in 3% (v/v) acetonitrile/0.1% (v/v) trifluoroacetic acid and 1 µg was directly injected onto a C18 1.7 µm, 130 Å, 75 µm X 250 mm M-class column (Waters) using a Thermo Ultimate 3000 RSLCnano UPLC. Peptides were eluted at 300 nL/minute using a gradient from 2% to 20% acetonitrile over 100 min into a Q-Exactive HF-X mass spectrometer (Thermo Scientific). Precursor mass spectra (MS1) were acquired at a resolution of 120,000 from 350 to 1550 m/z with an automatic gain control (AGC) target of 3E6 and a maximum injection time of 50 milliseconds. Precursor peptide ion isolation width for MS2 fragment scans was 1.4 m/z, and the top 12 most intense ions were sequenced. All MS2 spectra were acquired at a resolution of 15,000 with higher energy collision dissociation (HCD) at 27% normalized collision energy. An AGC target of 1E5 and 100 milliseconds maximum injection time was used. Raw files were searched against the Uniprot *Escherichia coli* databases UP000002032 and UP000000625 using Maxquant version 2.0.3.0 with cysteine carbamidomethylation as a fixed modification. Methionine oxidation and protein N-terminal acetylation were searched as variable modifications. All peptides and proteins were thresholded at a 1% false discovery rate (FDR). Statistical analysis was performed on log_2_-transformed iBAQ and LFQ intensities using limma (80).

### Plasmid-based P*cys-gfp* reporter system

To construct P*cys-gfp* reporter plasmids, the backbone of the low copy number (∼5 copies) plasmid pTHSSe_43 (82), including the pSC101 origin of replication with *repA* variant (E93V) gene and ampicillin resistance cassette, was linearized and amplified by PCR. A 312 bp fragment including the CysB-activated promoter of the *cysDNC* operon (P*cys*) through the start codon was amplified by PCR from gDNA from the ancestral *proBA*\*-*rpoS* relocation strain. A 777 bp fragment of the monomeric superfolder *gfp* gene lacking a start codon was amplified from pAA002. For the plasmid containing *cysB*, a 1,402 bp fragment including *cysB* with ∼200 bp upstream and downstream was amplified by PCR from gDNA from the ancestral strain. All primers used to amplify PCR fragments had ∼20 bp overlapping ends for Gibson assembly (83). Fragments were mixed together at an insert to vector ratio of approximately 4:1 with Gibson Assembly Master Mix (New England Biolabs) and incubated at 50 °C for 1 h. Gibson Assembly reactions were diluted 4-fold with sterile deionized water and the product was transformed into chemically competent *E. coli* DH5ɑ by heat shock at 42 °C for 30 sec according to manufacturer’s protocol. Following heat shock, cells were inoculated into SOC (BD Difco™) and allowed to recover at 37°C with shaking at 200 rpm for 1-2 h, then spread onto LB ampicillin plates. Ampicillin-resistant transformant colonies were screened for proper plasmid assembly by colony PCR. P*cys-gfp* reporter plasmids were purified using the Monarch^®^ Plasmid Miniprep Kit (New England Biolabs) according to the manufacturer’s protocol. The ancestral *proBA*\*-*rpoS* relocation *E. coli* strain was grown at 37 °C in LB to an OD_600_ of ∼0.3-0.5, washed with room temperature deionized water and immediately electroporated with 20-40 ng of purified plasmid. Following electroporation, cells were inoculated into SOC and allowed to recover at 37 °C with shaking at 200 rpm for ∼1 h, then spread onto LB ampicillin plates. Ampicillin-resistant and GFP-positive colonies were screened by colony PCR to confirm presence of the proper P*cys-gfp* reporter plasmid. All primers used to construct strains and plasmids are listed in *SI Appendix* Table S4.

Starter cultures were prepared by inoculating 5 μL of thawed frozen stock of each strain into 4 mL M9 supplemented with 0.2% glucose, 0.5 mM L-arginine, 0.4 mM L-proline, 20 μg/mL kanamycin and 150 μg/mL ampicillin and incubating cultures overnight at 37 °C with shaking at 200 rpm. The next day, 40 μL of each overnight culture was used to inoculate 4 mL of fresh M9 medium (1:100 dilution) supplemented as described above but with 100 μg/mL ampicillin. Cultures were grown at 37 °C with shaking at 200 rpm for ∼6 hours to an OD_600_ of ∼0.4-0.5. One mL of each culture was harvested by centrifugation at 8,000 x g for 3 min at room temperature. The cell pellets were resuspended in 1 mL of fresh M9 medium lacking carbon sources and antibiotics. A 100 μL aliquot of samples from both strains (six biological replicates and two technical replicates) was transferred to the wells of a 96-well Corning^®^ Costar black/clear bottom microplate. Sterile M9 medium was added to wells of the same plate as blank controls. Cell density (OD_600_) and GFP fluorescence (Excitation: 479 nm; Emission: 520 nm; Optics Position: Top) of each sample was measured using a BioTek Synergy H1 microplate reader (Agilent Technologies, Santa Clara, CA). The OD_600_ and GFP fluorescence of blank medium only controls were averaged and subtracted from all sample measurements as background. GFP fluorescence measurements for each sample were normalized by dividing by OD_600_ values. The normalized GFP fluorescence measurements for each set of technical replicates were averaged.

