## Supplementary material for "The transcriptomic and proteomic ramifications of segmental amplification": SI Appendix

**This PDF file includes**
Fig. S1. Population 2 contains two clades with distinct amplifications.
Fig. S2. The majority of cells in population 3 have an amplification nested inside a duplication.
Fig. S3. Relationship between gene copy number and expression at the RNA and protein levels.
Fig. S4. Population 8 evolved isolates have the same segmental amplification present at 6 copies.
Table S1. Amplicons present in evolved *proBA*-rpoS* populations.
Table S2. *E. coli* strains used in this work.
Table S3. Plasmids used in this work.
Table S4. Primers used in this work.

**Other supplementary materials for this manuscript include:**
Dataset S1. mRNA and protein fold change data for amplified genes in populations 3 and 8
Dataset S2. Proteomics data and differential expression analysis for populations 3 and 8
Dataset S3. RNA-sequencing transcriptomic data and differential expression analysis for all genes in populations 3 and 8

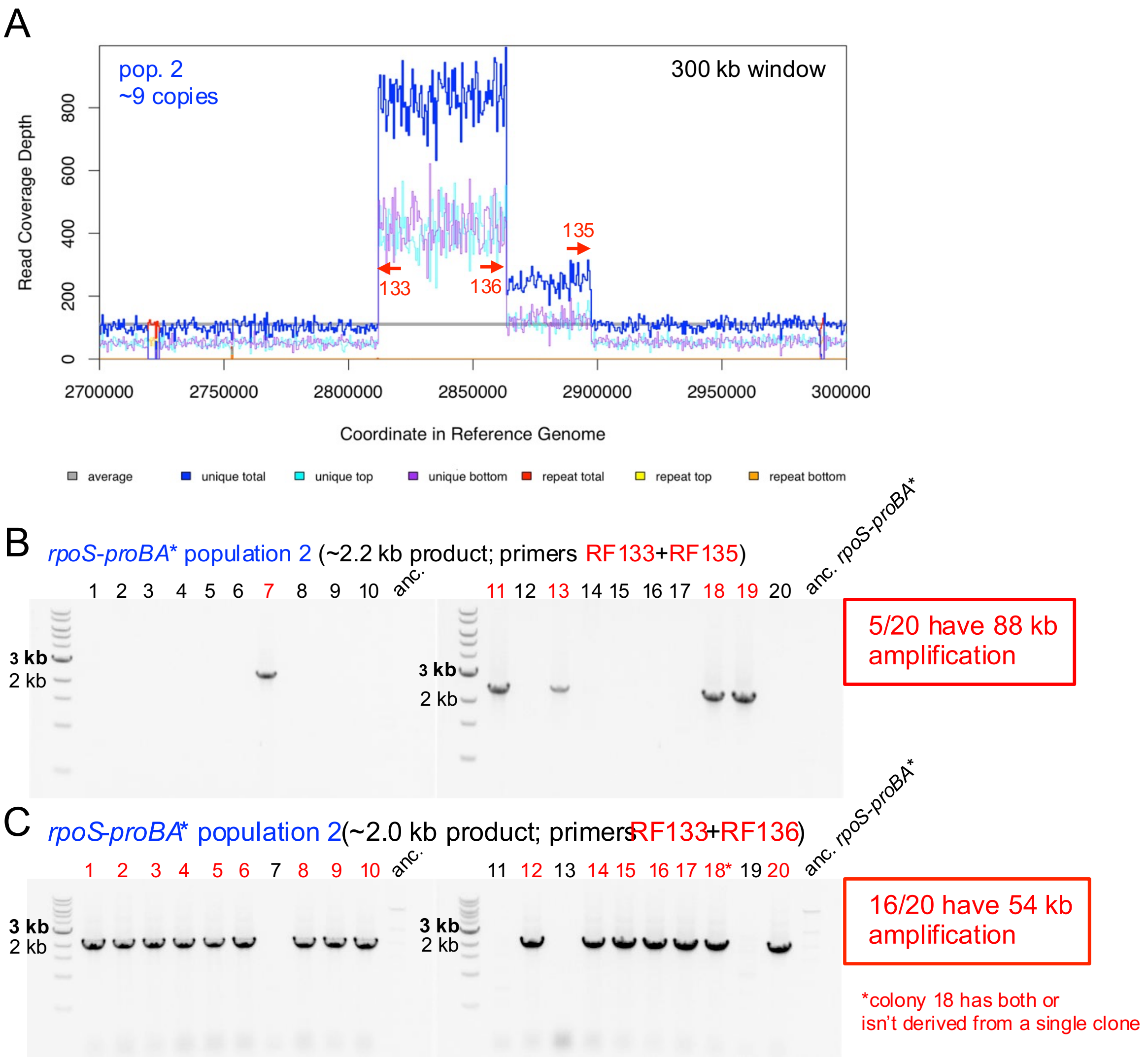


**Fig. S1. Population 2 contains two clades with distinct amplifications.** (A) Read coverage map for population 2 shows 88 and 54 kb amplifications. Red arrows indicate binding sites for primers used to interrogate potential amplification junctions. (B) Colony PCR results for 20 evolved isolates using primers RF133 and RF135. (C) Colony PCR results for 20 evolved isolates using primers RF133 and RF136.

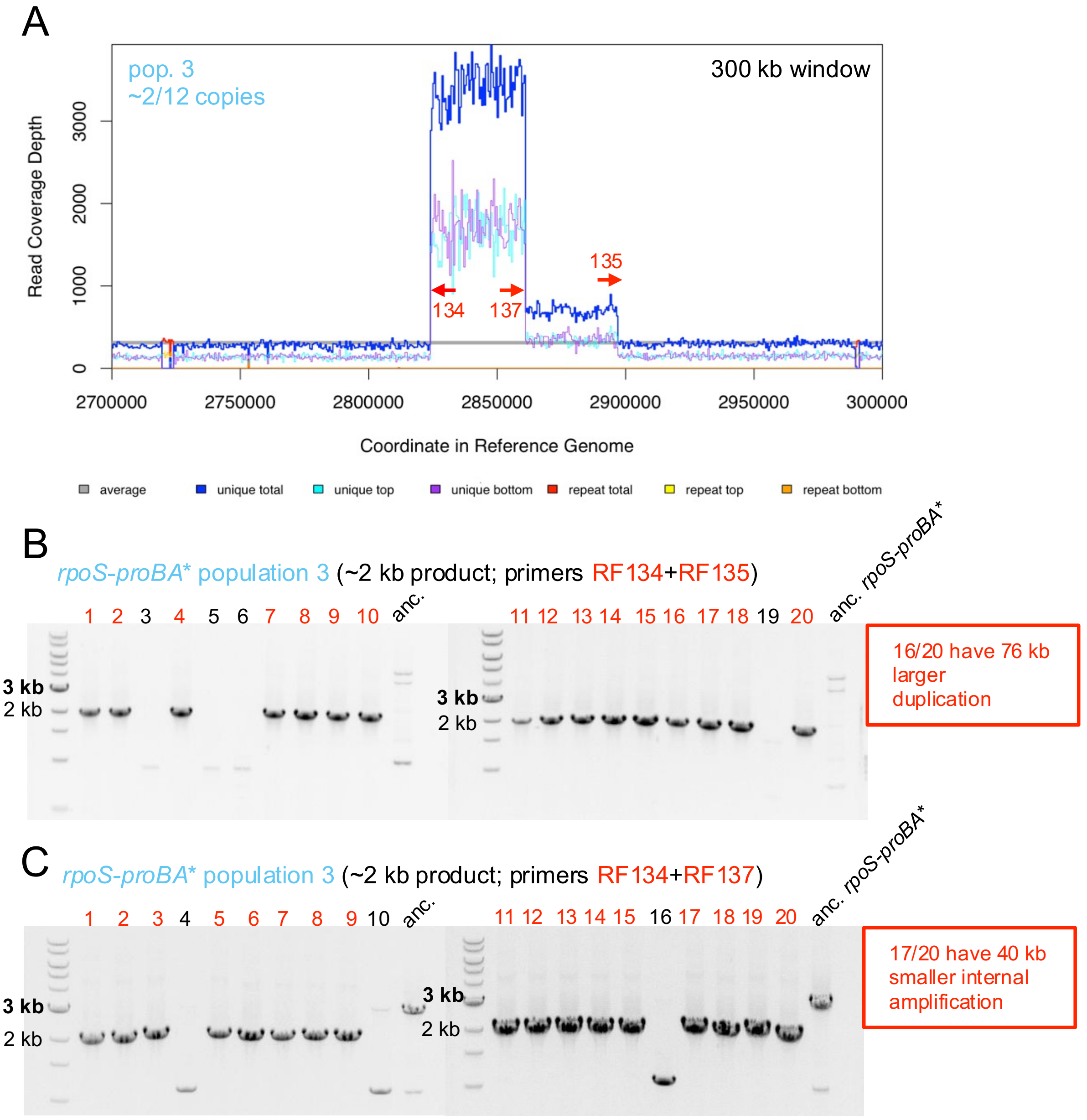


**Fig. S2. The majority of cells in population 3 have an amplification nested inside a duplication.** (A) Read coverage map for population 3 shows 76 and 40 kb amplifications. Red arrows indicate binding sites for primers used to interrogate potential amplification junctions. (B) Colony PCR results for 20 evolved isolates using primers RF134 and RF135. (C) Colony PCR results for 20 evolved isolates using primers RF134 and RF137.

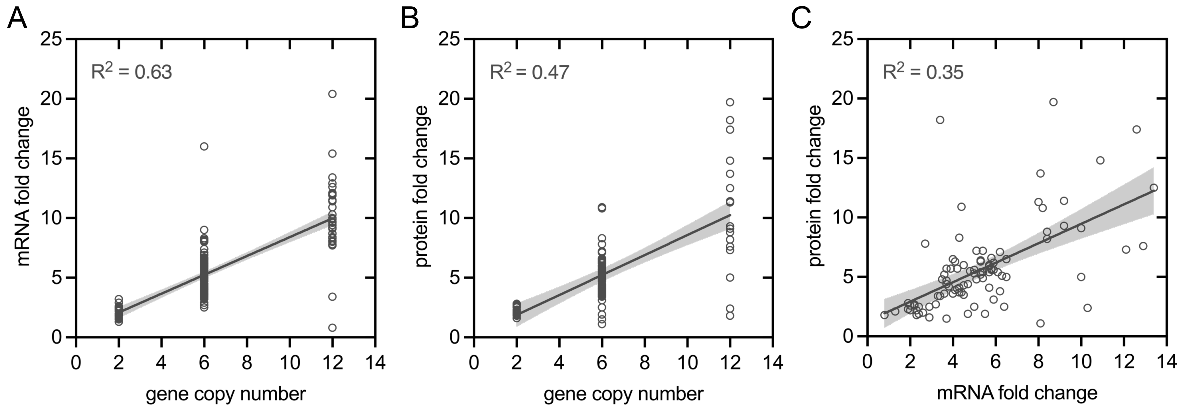


**Fig. S3. Relationship between gene copy number and expression at the RNA and protein levels.** (A) Correlation between mRNA fold change and gene copy number. (B) Correlation between protein fold change and gene copy number. (C) Correlation between protein fold change and mRNA fold change. $R^2$ values were calculated by simple linear regression. Shaded areas indicate 95% confidence intervals. Data points for *hycB* (<5 read counts/replicate) and HycE (<1 peptide/replicate) were removed from graphs and not included in linear regression analysis because their levels in the reference strain were very low and thus their fold-changes were not reliable.

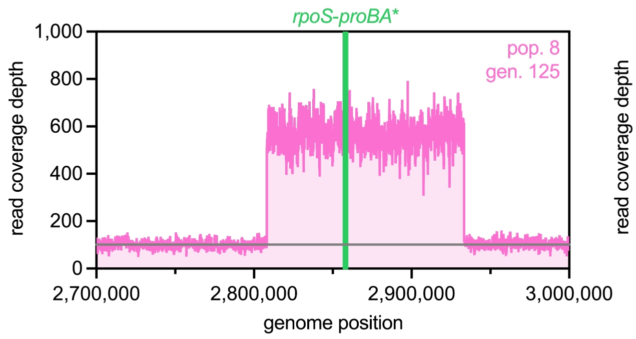


**Fig. S4. Population 8 evolved isolates have the same segmental amplification present at 6 copies.** Read coverage maps for population 8 isolates from generation 125 (left) and generation 186 (right) showing segmental amplifications.

**Table S1. Amplicons present in evolved *proBA*-rpoS* populations.**

| pop. | gen. | upstream junction position[a] (gene) | downstream junction position[a] (gene) | IS element involved | amplicon size (kb) | *proA** copy number |
|---|---|---|---|---|---|---|
| 1 | 155 | 2808170 (*serV*) | 2864360 (*ispD*) | *insL1* (IS186) | 56.2 | 7.8 |
| | | | 2896344 (*ygcE/queE*) | | 88.2 | |
| 2 | 146 | 2808170 (*serV*) | 2862506 (*umpG*) | *insL1* (IS186) | 54.3 | 8.8 |
| | | | 2896283 (*ygcE/queE*) | | 88.1 | |
| 3 | 198 | 2820227 (*srlQ*) | 2859849 (*nlpD*) | *insAB1* (IS1) | 39.6 | 11.8 |
| | | | 2859931(*nlpD*) | | 39.7 | |
| | | | 2895963 (*ygcE/queE*) | | 75.7 | |
| 4 | 182 | 2851868 (*ygbJ*) | 2860822 (*nlpD*) | none | 9.0 | 7.7 |
| | | 2808170 (*serV*) | 2896344 (*ygcE/queE*) | *insL1* (IS186) | 88.2 | |
| 5 | 157 | 2808170 (*serV*) | 2861262 (*pcm*) | *insL1* (IS186) | 53.1 | 7.2 |
| 6 | 159 | 2808170 (*serV*) | 2896283 (*ygcE/queE*) | *insL1* (IS186) | 88.1 | 7.0 |
| 7 | 172 | 2835653 (*hycE*) | 2868717 (*cysD/iap*) | *insAB1* (IS1) | 33.1 | 8.5 |
| | | 2808170 (*serV*) | 2896283 (*ygcE/queE*) | *insL1* (IS186) | 88.1 | |
| 8 | 186 | 2808170 (*serV*) | 2874835 (*casB*) | none | 66.7 | 5.8 |
| | | | 2933283 (*rlmM*) | *insL1* (IS186) | 125.1 | |

[a]Position numbers based on reference genome for ancestral *proBA*-rpoS* strain (RF90).

**Table S2. *E. coli* strains used in this work.**

| Strain | Genotype/description | Source/reference |
|---|---|---|
| JW3930 (AM008) | Δ*argC*::$kan^R$ Keio collection strain; derived from *E. coli* BW25113 | (1) |
| AM187 | -45 C > T *proB* promoter mutation (M2), *proA** (E383A), +*yfp* inserted 282 bp downstream *proA**, Δ*fimAICDFGH*, Δ*csgBAC;* derived from JW3930/AM008 | (2, 3); Genbank accession number CP037857.2; M2 promoter mutation described in (4) |
| AM327 | Δ82 bp in *rph* 90 bp upstream of *pyrE*, derived from AM187 | (2) |
| RF70 | Δ*proBA*-yfp*::$sacB\text{-}gtm^R$; derived from AM327 | this study |
| RF72 | Δ*proBA*-yfp;* derived from RF70 | this study |
| RF83 | Δ*proBA*-yfp*, *ygbN-tetR-ccdB-cat-rpoS*; derived from RF72 | this study |
| RF90 | Δ*proBA*-yfp,* M2-*proBA*-rpoS;* derived from RF83; ancestral strain for laboratory evolution | this study |

**Table S3. Plasmids used in this work.**

| Plasmid | Description | Source/reference |
|---|---|---|
| pJQ200SK | suicide vector containing sucrose-inducible *sacB* and constitutive $gtm^R$ selection-counterselection cassette | (5) |
| pDLM3 | plasmid containing anhydrotetracycline-inducible *tetR-ccdB-cat* selection-counterselection cassette | (6) |
| pSIM6 | temperature-sensitive plasmid containing heat-inducible λ-Red genes (*exo*, *bet*, *gam*) for recombineering; $amp^R$ | (7) |
| pTHSSe_43 | Low-copy-number (~5 copies) plasmid with pSC101 origin and encoding RepA E93V variant; $amp^R$ | (8) |
| pET-His6-msfGFP-TEV | plasmid encoding monomeric superfolder green fluorescence protein (msfGFP); $amp^R$ | Addgene plasmid # 48287 (9) |
| pAA002 | plasmid encoding ProA-msfGFP fusion protein derived from pET-His6-msfGFP-TEV (9); $amp^R$ | (10) |
| pTHSSe-Pcys-msfgfp | plasmid containing reporter system consisting of msfGFP under control of CysB-activated Pcys promoter but lacking *cysB* gene; pTHSSe_43 vector backbone; $amp^R$ | This study |
| pTHSSe-Pcys-msfgfp-cysB | plasmid containing reporter system consisting of msfGFP under control of CysB-activated Pcys promoter and including *cysB* gene; pTHSSe_43 vector backbone; $amp^R$ | This study |

**Table S4. Primers used in this work.**

| Name | Sequence (5' → 3')[a] | Use |
|---|---|---|
| RF35-fwd | CATTACGGATTCACTGGCCG | amplifying *tetR-ccdB-cat* cassette |
| RF36-rvs | CGGTAAATAGCTTGCCTGCTC | amplifying *tetR-ccdB-cat* cassette |
| RF61-fwd | CGATCGACTCTAGCTAGAGGATC | amplifying *sacB-gtm*$^R$ cassette |
| RF62-rvs | GAAACGGATGAAGGCACGAAC | amplifying *sacB-gtm*$^R$ cassette |
| RF71-fwd | GGCGTCATTTTGCGTGTTTCAG | amplifying region upstream of *proBA* |
| R72-rvs | cctctagctagagtcgatcgCAACCATCTGCG AGCAAAGC | amplifying region upstream of *proBA* |
| RF73-fwd | ttcgtgccttcatccgtttcGGCATGGACGAGC TGTACAAG | amplifying region downstream of *proBA* |
| RF74-rvs | CCACTCAGCGTTAATTACCTACACG | amplifying region downstream of *proBA* |
| RF81-rvs | ttgtacagctcgtccatgccCAACCATCTGCGA GCAAAGC | amplifying region upstream of *proBA* to replace *sacB-gtm*$^R$ with markerless deletion of *proBA* |
| RF82-fwd | gctttgctcgcagatggttgGGCATGGACGAG CTGTACAAG | amplifying region upstream of *proBA* to replace *sacB-gtm*$^R$ with markerless deletion of *proBA* |
| RF88-fwd | GCAACAAAACGCCATGCTTT | amplifying *proBA** with mutant M2 promoter |
| RF89-rvs | TGGCCTTGTGAATTATAAACGC | amplifying *proBA** with mutant M2 promoter |
| RF91-fwd | GTCGGCTTTATTATTCTTGCGCC | amplifying region upstream of the target site between *rpoS* and *ygbN* |
| RF97-rvs | cggccagtgaatccgtaatgGGCTGGCCTTTT CTGTGCAC | amplifying region upstream of the target site between *rpoS* and *ygbN* |
| RF94-fwd | GAAGATGCGGAATTTGATGAGAACG | amplifying region downstream of the target site between *rpoS* and *ygbN* |
| RF98-rvs | gagcaggcaagctatttaccgTCGCTTGAGAC TGGCCTTTCTG | amplifying region downstream of the target site between *rpoS* and *ygbN* |
| RF99-rvs | aaagcatggcgttttgttgcGGCTGGCCTTTTC TGTGCAC | amplifying region upstream of the target site between *rpoS* and *ygbN* to replace *tet*$^R$-*ccdB-cat* with M2-*proBA** |
| RF100-fwd | cgtttataattcacaaggccaTCGCTTGAGACT GGCCTTTCTG | amplifying region downstream of the target site between *rpoS* and *ygbN* to replace *tet*$^R$-*ccdB-cat* with M2-*proBA** |
| RF133-rvs | CCGCGTCTCATCTTTATCGGTG | checking presence of 54/88 kb amplification in evolved clones from population 2 (binds near upstream junction) |
| RF134-rvs | GGCGCTATCAGTAATCGCAGC | checking presence of 40/76 kb amplification in evolved clones from population 3 (binds near upstream junction) |
| RF135-fwd | TCGTATGAAGTGATGCATGGGG | checking presence of 88 kb or 76 kb amplification in evolved clones from populations 2 or 3, respectively (binds near downstream junction) |
| RF136-fwd | CGTCACGCGAATACCTTTGATTTG | checking presence of 54 kb amplification in evolved clones from population 2 (binds near downstream junction) |
| RF137-fwd | CACCAGGTTGCGTATGTTGAGAAG | checking presence of 40 kb amplification in evolved clones from population 3 (binds near downstream junction) |
| RF138-fwd | GCAAACGTGGTAGAGAAGGCAC | amplifying *prlF-yhaV* |
| RF139-rvs | GTCAGTCGAGCCATCAGTGGAAG | amplifying *prlF-yhaV* |
| RF140-fwd | GGCTGTAAAAGGACAGTGAATCATG | sequencing *prlF* |
| RF141-rvs | GCTGAATACGGTATAGGCATCTG | sequencing *prlF* |

| | | |
|---|---|---|
| RF178-fwd | tatcacgaggGAATAAAGCATACGCCGAG | amplifying promoter of *cysDNC* operon (P*cys*) including its start codon |
| RF179-rvs | ccttgctcacCATAACCGTTCCTTTGCAATAC | amplifying promoter of *cysDNC* operon (P*cys*) including its start codon |
| RF180-fwd | aacggttatgGTGAGCAAGGGCGAGGAG | amplifying *msf*-*gfp* lacking start codon for plasmids with and without *cysB* |
| RF181-rvs | tgctttattcCCTCGTGATACGCCTATTTTTATAG | linearizing and amplifying pTHSSe_43 vector backbone for plasmids with and without *cysB* |
| RF182-fwd | aacctgacagGAGCGGTATCAGCTCACTC | linearizing and amplifying pTHSSe_43 vector backbone for plasmid containing *cysB* |
| RF183-rvs | aattcgctggGCCACCTGACGTCTAAGAAAC | amplifying *msf*-*gfp* lacking start codon for plasmid containing *cysB* |
| RF184-fwd | gtcaggtggcCCAGCGAATTTACGAATC | amplifying *cysB* |
| RF185-rvs | gataccgctcCTGTCAGGTTTTTATAAACAAAG | amplifying *cysB* |
| RF186-rvs | gataccgctcGCCACCTGACGTCTAAGAAAC | amplifying *msf*-*gfp* lacking start codon for plasmid lacking *cysB* |
| RF187-fwd | gtcaggtggcGAGCGGTATCAGCTCACTC | linearizing and amplifying pTHSSe_43 vector backbone for plasmid lacking *cysB* |
| 1357-fwd | CGTGCAGGCGATTGATAA | quantifying *proA** copy number by qPCR |
| 1358-rvs | CTGTTCACGGCACAGTTT | quantifying *proA** copy number by qPCR |
| 1359-fwd | CGTAGATCTGACGGTGAATTT | quantifying *gyrB* copy number by qPCR |
| 1360-rvs | CGTTGGTGTTTCGGTAGTA | quantifying *gyrB* copy number by qPCR |
| 1361-fwd | CCCGTGGCTGAAAGTTAAA | quantifying *icd* copy number by qPCR |
| 1362-rvs | CAGGTTCATACAGGCGATAAC | quantifying *icd* copy number by qPCR |

[a]lower case letters indicate non-annealing sequence overlaps used for splicing by overlap extension PCR or Gibson assembly.
